# Dimensionality and ramping: Signatures of sentence integration in the dynamics of brains and deep language models

**DOI:** 10.1101/2023.02.28.530443

**Authors:** Théo Desbordes, Yair Lakretz, Valérie Chanoine, Maxime Oquab, Jean-Michel Badier, Agnès Trébuchon, Romain Carron, Christian-G. Bénar, Stanislas Dehaene, Jean-Rémi King

## Abstract

A sentence is more than the sum of its words: its meaning depends on how they combine with one another. The brain mechanisms underlying such semantic composition remain poorly understood. To shed light on the neural vector code underlying semantic composition, we introduce two hypotheses: First, the intrinsic dimensionality of the space of neural representations should increase as a sentence unfolds, paralleling the growing complexity of its semantic representation, and second, this progressive integration should be reflected in ramping and sentence-final signals. To test these predictions, we designed a dataset of closely matched normal and Jabberwocky sentences (composed of meaningless pseudo words) and displayed them to deep language models and to 11 human participants (5 men and 6 women) monitored with simultaneous magneto-encephalography and intracranial electro-encephalography. In both deep language models and electrophysiological data, we found that representational dimensionality was higher for meaningful sentences than Jabberwocky. Furthermore, multivariate decoding of normal versus Jabberwocky confirmed three dynamic patterns: (i) a phasic pattern following each word, peaking in temporal and parietal areas, (ii) a ramping pattern, characteristic of bilateral inferior and middle frontal gyri, and (iii) a sentence-final pattern in left superior frontal gyrus and right orbitofrontal cortex. These results provide a first glimpse into the neural geometry of semantic integration and constrain the search for a neural code of linguistic composition.

**Significance statement:** Starting from general linguistic concepts, we make two sets of predictions in neural signals evoked by reading multi-word sentences. First, the intrinsic dimensionality of the representation should grow with additional meaningful words. Second, the neural dynamics should exhibit signatures of encoding, maintaining, and resolving semantic composition. We successfully validated these hypotheses in deep Neural Language Models, artificial neural networks trained on text and performing very well on many Natural Language Processing tasks. Then, using a unique combination of magnetoencephalography and intracranial electrodes, we recorded high-resolution brain data from human participants while they read a controlled set of sentences. Time-resolved dimensionality analysis showed increasing dimensionality with meaning, and multivariate decoding allowed us to isolate the three dynamical patterns we had hypothesized.

## Introduction

During sentence comprehension, the human brain must bind successive words into an integrated representation. The neural basis of such compositionality has been partially studied for two word phrases (Pylkkänen, 2019, 2020), but remains largely unknown for longer constituents. Here, combining ideas from the neural population framework (Georgopoulos et al., 1986; Yuste, 2015; Ebitz & Hayden, 2021) and vector models of linguistic composition (Smolensky, 1990), we hypothesize that the neural manifold representing sentences should grow with the progressive addition of meaningful words. Two sets of predictions unfold from this vector framework (Figure 1).

**Figure 1.**
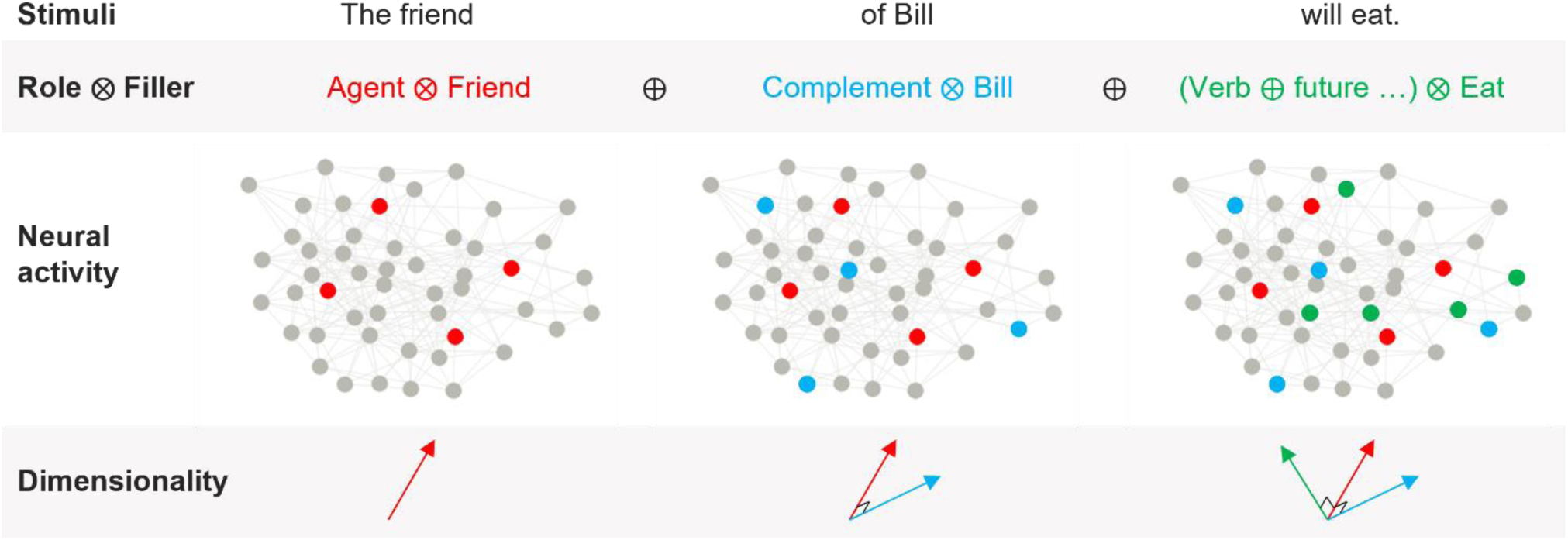
Logic of the research As successive words are integrated into a sentence-level representation, a variety of models predict that they are integrated into an internal representation which combines word identity with syntactic role, and sums the resulting filler-role bindings (Smolensky, 1990). Assuming that these neural vectors are sparse, this schema makes two predictions. First, average neural activity, as indexed for instance by high-gamma activity, should increase; and second, the dimensionality of the neural manifold evoked by a variety of such sentences should increase, because across a variety of words, the vectors point in a variety of directions and therefore span a larger space. Such increases would not be expected (or to a much extent) for a list of meaningless consonant strings or for a Jabberwocky sentence -- the latter being constantly represented as {agent:meaningless} + {complement:meaningless} + {verb:meaningless}…

First, for each word that the subject successfully integrates, we predict an increase in intrinsic dimensionality of the corresponding neural representation, i.e. the number of independent dimensions that participate in the encoding (Carreira-Perpinán, 1997), or “the dimensionality of the manifold that approximately embeds the data” (Del Giudice, 2021). Intrinsic dimensionality should increase as new meaning elements are put together because each word, once bound to its syntactic role, should occupy a distinct axis in neural vector space (Smolensky, 1990) : when we combine real words with one another, we generate a meaning which is more than the sum of its parts, and thus requires additional dimensions. Intuitively, one can think of concept cells (Quiroga et al., 2005) being recruited to encode incoming elements and their relationships. For Jabberwocky sentences, where meaningless pseudowords replace actual words (Mazoyer et al., 1993; Hahne & Jescheniak, 2001), we predict the recruitment of a reduced set of semantic dimensions because pseudowords carry little or no meaningful information.

Our second set of predictions relates to the dynamics of composition. We predict that meaningful composition will lead to a growing superposition of neural codes each recruiting additional dimensions, and thus leading to ramping neural activity over the course of the sentence (Figure 1). Several studies support this prediction (Bastiaansen et al., 2009; Pallier et al., 2011; Fedorenko et al., 2016; Nelson et al., 2017), but they did not clearly separate multi-word integration from other semantic processes of lexical access and sentence wrap-up. Here, we used multivariate decoding with temporal generalization (King & Dehaene, 2014; Fyshe, 2020) to disentangle them. We derived theoretical generalization matrices for the predicted dynamics of classifiers trained to separate normal and Jabberwocky sentences for each process (Figure 2D). Those three stages are not tied to a particular theory of sentence processing (Just & Carpenter, 1980; Seidenberg & McClelland, 1989; Steedman, 2001; Lewis & Vasishth, 2005); rather, we propose them as necessary steps in sentence comprehension.

**Figure 2:**
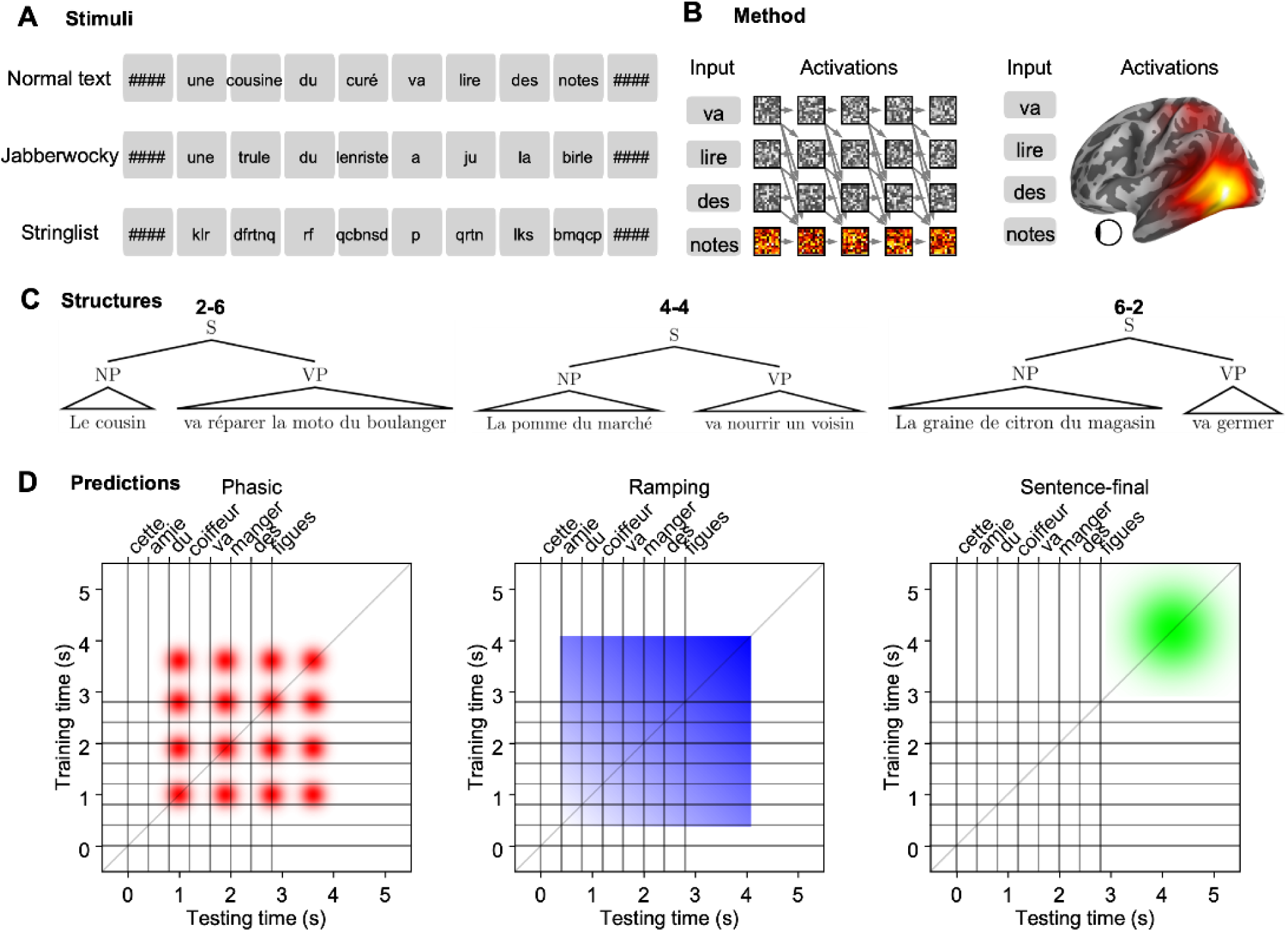
Experimental design A. Example stimuli for the 3 conditions: normal text, jabberwocky, and list of consonant strings (stringlist). Masks of ’#’ were presented before and after each sentence in order to keep visual masking approximately constant. B. Extraction of activations from Neural Language Models and combined MEG-sEEG brain recordings. C. Example of the 3 syntactic structures used in this study, varying in the relative size of the NP and the VP. D. Theoretical temporal generalization patterns. Regions involved in lexical processing of single words would differentiate normal content words from jabberwocky content words based on lexical access mechanisms (the function words are the same in both conditions), yielding a phasic pattern. Regions involved in compositional processes should exhibit a ramping generalization pattern, where normal and jabberwocky sentences become more and more differentiated with each incoming word. Finally, regions involved in wrap-up processes would separate normal and jabberwocky only after the sentence is finished, leading to a sentence-final pattern. See Figure 2-1 for electrode placement.

First, lexical access is predicted to elicit a phasic, transient response that separates normal words from Jabberwocky (Just & Carpenter, 1980; Caramazza, 1997) (Figure 2D, red), especially in the anterior fusiform gyrus (FuG) and superior and middle temporal gyri (STG, MTG) (Nobre et al., 1994; Binder et al., 2003; Woolnough et al., 2020).

Second, multi-word integration is predicted to elicit ramping dynamics, characterized by an increasingly strong square pattern in the temporal generalization matrix (Figure 2D, blue). This pattern is predominantly expected in Inferior Frontal Gyrus (IFG) and part of superior temporal sulcus (STS) (Pallier et al., 2011; Fedorenko et al., 2016; Nelson et al., 2017).

Last, sentence-final wrap-up processes are predicted to occur after the last word and elicit a sentence-final separation of normal versus Jabberwocky sentences (Figure 2D, green). Indeed, readers are known to pause at the end of sentences to integrate, interpret, and incorporate the constituting elements into the general context of the discourse (Just & Carpenter, 1980). Behavioral eye-tracking studies have evidenced an increased reading time for sentence-final words (Warren et al., 2009; Kuperman et al., 2010). End-of-sentence neural signatures of in-sentence grammatical gender violation (Molinaro et al., 2008) and syntactic complexity (Pattamadilok et al., 2016) have also been observed.

We tested the above predictions in both humans and artificial neural networks. Convergent findings would support the view that those properties are general features of linguistic composition.

## Methods

### Ethics

Eleven right-handed individuals (5 men and 6 women; age range=25–57, mean= 40, SD= 9.4) with intracranial stereotactic electrodes implantation as part of their treatment for refractory epilepsy gave their informed consent to participate in our study, in accordance with the ethic evaluation RCB 2018-A02363-52. All patients were implanted with depth electrodes for clinical purpose (presurgical evaluation) in the Epileptology and Cerebral Rythmology Department of the Timone Hospital (Marseille, France). Neuropsychological assessment indicated that all patients had intact language functions. Their reading ability was controlled by means of a French version of a reading test (test Malabi, © Unité INSERM-CEA de Neuroimagerie Cognitive).

### Stimuli and task

8-word-long sentences were presented to the participants in a Rapid Stream Visual Presentation with an SOA of 400ms. Each sentence was preceded and followed by visual masks (####) in order to keep forward and backward masking constant (Figure 2A).

The stimuli were generated using a custom sentence generator script that constructs a wide range of sentences from a finite vocabulary set, respecting several constraints. (a) All sentences were 8 words long. (b) Each sentence comprised a systematic alternation of short function words (determiners and auxiliary) and longer content words (nouns and verbs). (c) The sentences consisted of a Noun Phrase (NP) followed by a Verb Phrase (VP). The NP consisted of a determiner, a noun (the subject of the sentence) and optionally one or two prepositions. The VP consisted of an auxiliary, a verb and optionally a determiner and a noun (the object of the verb) and one or two prepositions. (d) There were 3 possible syntactic structures that varied in the size of the NP and VP. They could both be of size 2, 4, or 6 words, while their sum was always equal to 8 words (Figure 2C).

The vocabulary consisted of 9 determiners, i.e., 3 for each gender in the singular form and 3 for the plural form; 10 verbs that could appear in singular or plural, in the present or past tense (40 different forms); and 75 nouns, among those 46 could appear in the singular or plural form (the others were always singular), 46 were masculine (26 were used as subjects and objects, the rest were appeared in prepositional phrases), 26 were feminine (all were used as subjects and objects) and 3 could appear in either masculine or feminine form (used as subjects only).

The total number of distinct sentences the script could generate was 791,754. For each subject, in the normal and jabberwocky conditions, we sampled an equal number of each syntactic structure, as well as an equal number of feminine/masculine and singular/plural subjects and objects and an equal number of present/past tense for the verb.

The Jabberwocky stimuli were designed by hand, by changing one or two letters of each content word to create a corresponding nonword, while keeping the morphological markers present.

Strings lists consisted of strings of consonants with the same length as the original words. These letter strings did not have any morphosyntactic information and thus constituted a low-level, mainly visual, control to the linguistic stimuli used in the experiment.

The task was for the participant to detect the presence of target words in the sentences and press the response button as fast as possible when the target was present. The target was present in 1/11 sentences. Participants performed 330 trials in total (cut down in 6 blocks), composed of 300 non-target trials (120 normal sentences, 120 jabberwocky sentences, and 60 strings of consonants) and 30 target sentences of a randomly selected condition containing the target words, that were discarded from the analyses.

### sEEG and MEG data acquisition and preprocessing

MEG and SEEG recordings took place simultaneously in a dimly illuminated, magnetically shielded room. Recordings were obtained from subjects in supine position to limit the movement during the recording.

MEG signals were acquired with a 248-channel biomagnetometer system (Magnetometers. 4D Neuroimaging, San Diego, CA, USA located in the MEG facility, Timone Hospital, Marseille). The data were recorded continuously with a band- DC-800 Hz bandwidth with a sampling rate of 2034.51 Hz.

SEEG recordings were performed using intracerebral multiple contact electrodes (10–15 contacts, length: 2 mm, diameter: 0.8mm, 1.5 mm apart from edge to edge) placed intracranially according to Talairach’s stereotactic method (Bancaud et al., 1970; Talairach et al., 1992).

SEEG as well as EOG and ECG signals, to facilitate the ulterior rejection of eye movements, blinks, and cardiac artifacts, were simultaneously recorded with MEG (Badier et al., 2017) with a band- 0.01-1000 Hz bandwidth with a sampling rate of 2500 Hz using a 256-channel BrainAmp amplifier system (Brain Products GmbH, Munich, Germany). SEEG was then interpolated at the sampling rate of the MEG thanks to triggers.

In order to determine the location of the head with respect to the MEG helmet, five coils were fixed on the subject’s head. The position of these coils as well as the surface of the head were digitized with a 3-D digitizer (Polhemus Fastrack, Polhemus Corporation, Colchester, VT, USA), and head position was measured at the beginning and at the end of each run. The head shape obtained from the digitization of the head was used to check and eventually compensate for differences in head position between runs or to match to the participant’s MRI. All stimuli were presented to the subjects on a mirror by a back-projection system where an LCD projector was placed outside the magnetically shielded room in order to avoid interfering electrical apparatus. The distance between the participant’s eyes and the screen on which stimuli were displayed was similar across patients. A trigger square invisible to the participant was projected onto a photodiode which was used to signal the presence of a stimulus on-screen and to synchronize the MEG and EOG/ECG recordings.

### Among the eleven patients, nine underwent an MEG recording at the same time

The sEEG and MEG data were band-pass filtered at 0.3-500hz, notch filtered at 50h and the first 3 harmonics to remove line noise. The data were then downsampled to 100hz and clipped at 10 times the standard deviation either side of the median value, separately for each channel. We then used an automatic detection procedure for bad channels in which the temporal variance is computed for each channel and a value above or below 25 times the median variance over channels leads to rejection.

Epochs were constructed keeping time points from -0.5 to 5.5 seconds after the onset of the first mask. We then used a procedure to reject bad epochs similar to the one we used for channels: the variance was now computed over time and the remaining channels, separately for each epoch, and a value above or below 5 times the median over epochs lead to rejection. Baseline correction was then applied using the 400ms interval between the onsets of the first mask and the first word. The data were then smoothed using a 100ms hanning window. Finally, Common Median Referencing was applied using all channels but the ones that were marked as bad. All the data preprocessing was done using the MNE-Python software (Gramfort et al., 2013).

The high-gamma activity (in Figure 3) was extracted using a Morlet transform and 8 frequency bands linearly spaced between 70 and 150 hertz, then combined with Principal Component Analysis, which we found to be more robust than simple averaging. For all subsequent analyses we used raw voltage instead high-gamma because we found the overall decoding performance to be better. However, we replicated the decoding and dimensionality results with high-gamma and did not find major differences.

**Figure 3:**
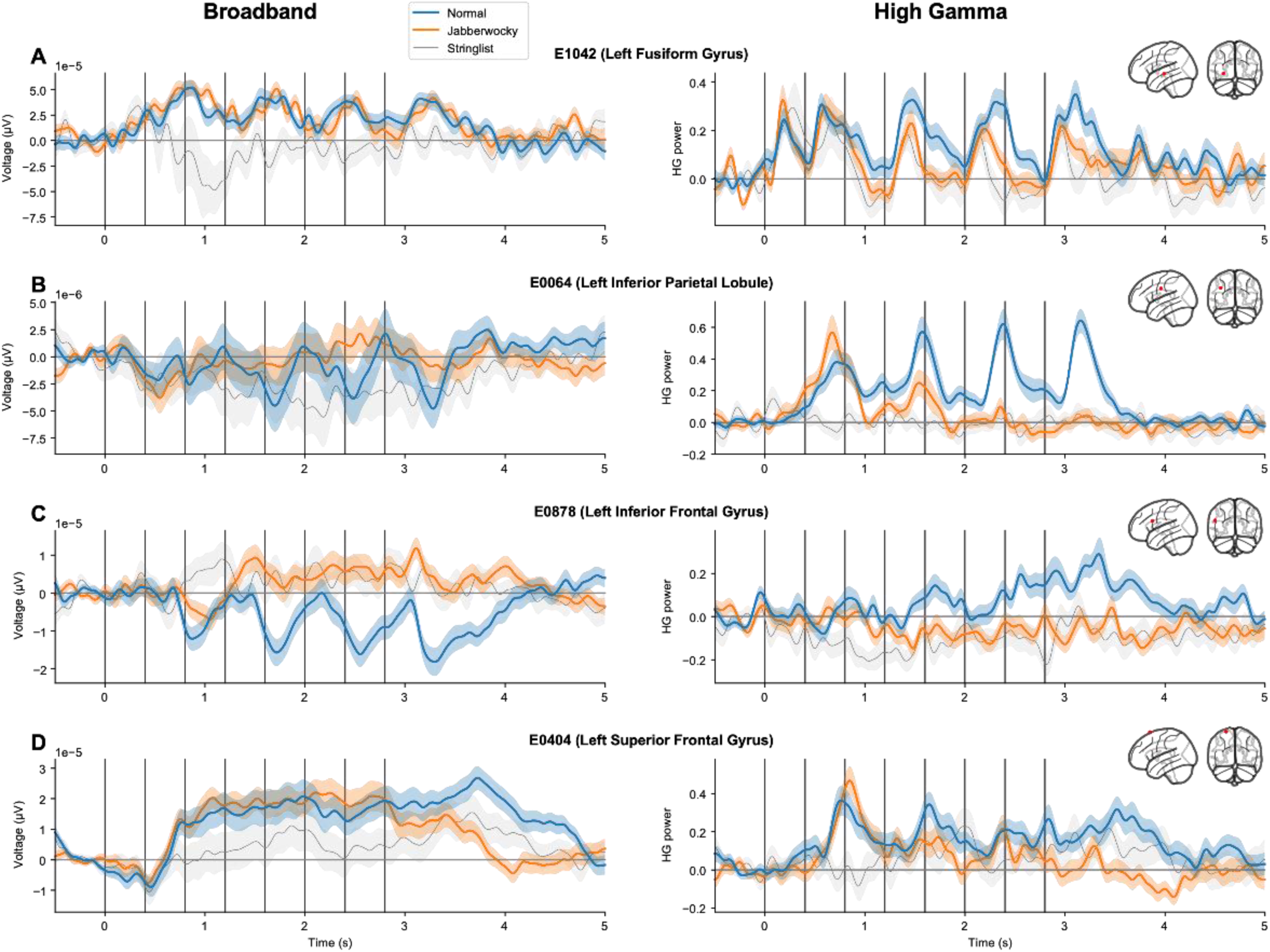
Illustrative profiles of human sEEG responses compatible with the postulated phasic, ramping and sentence-final patterns. Four examples of electrodes responding to normal sentences (blue), Jabberwocky sentences (orange), and string of consonants (gray). Each line shows the local field potential (Voltage, Left) and the high-gamma (HG power, Right) for the same electrode. The 8 vertical lines represent the onset of each word in the sentence. See Figure 3-1 for additional illustrative profiles of human sEEG responses. See Figure 3-2 for activations from the Transformer that exhibit similar dynamics.

### Localizer test

To boost the statistical power, we selected language-specific electrodes using a two-sample temporal cluster permutation test. Specifically, we tested whether the conditions (normal text, jabberwocky) and (stringlist) were different at the whole-epoch level. We kept electrodes that contained at least one cluster after the permutation test and FDR correction. We used 1000 permutations and threshold-free cluster enhancement (Smith & Nichols, 2009) with a starting threshold of 0 and a step of 0.1.

### Dimensionality Analysis

Within the large dimensionality of the overall neural space (equal to the number of relevant neurons), only a much smaller vector subspace is actually used for encoding (Ebitz & Hayden, 2021). To quantify this intrinsic dimensionality (ID), we used a previously reported method based on Principal Component Analysis (PCA) (Gao et al., 2017; Elmoznino & Bonner, 2022), sometimes called the participation ratio (Sorscher et al., 2022). This method quantifies intrinsic dimensionality as follows:

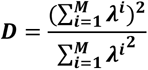

where λ^*i*^are the eigenvalues of the neural covariance matrix (i.e., the eigenvalues whose corresponding eigenvectors are the principal components of the dataset), and M is the number of channels (electrodes or magnetometers). This gives a continuous measure of the number of principal components needed to explain most of the variance in a dataset. Intuitively, one can check that if the data varies only along a single dimension, all of the variance will be explained by the first principal component, hence a single eigenvalue λ^1^ will be non-zero, and therefore the formula implies that D = 1. On the contrary, if the signals in all channels vary independently of each other (and with similar magnitude), such that each principal component explains an equal part of the variance, then D will be equal to the number of channels. Between those two extremes, D estimates the approximate number of dimensions that vary significantly in the brain signals.

To calculate D, we computed a PCA on 0.4 s sliding time windows, combining time points and trials (such that the PCA’s input is a n_times*n_trials by n_channels matrix), separately for each of the 3 conditions (normal, Jabber, stringlist), and computed the ID of the resulting eigenspectrum using the aforementioned formula. This analysis was repeated 10 times with different (non-overlapping) parts of the data to get an average D and the corresponding standard error bars.

This measure of intrinsic dimensionality has been used for some time in statistics and machine learning (Carreira-Perpinán, 1997; Campadelli et al., 2015) and has recently gained traction in the neuroscience domain, where low-dimensional intrinsic dynamics were found in high-dimensional neural recordings (Machens et al., 2010; Churchland et al., 2012; Mante et al., 2013; Xie et al., 2022). These low-dimensional dynamics have also been found to emerge in trained neural networks (Laje & Buonomano, 2013; Recanatesi et al., 2021) and have been assigned important roles in various theories of neural computation (Gallego et al., 2017; Gao et al., 2017; Vyas et al., 2020; Ebitz & Hayden, 2021; Sorscher et al., 2021).

Importantly, intrinsic dimensionality is not equivalent to overall neural activity. Rather, the two characteristics may vary independently: very high activity in a single dimension will bring a low dimensionality, whereas the same amount of activity distributed across more dimensions yields a high dimensionality. Furthermore, while overall neural activity is an average measure, dimensionality reflects the dispersion of activity over a set of trials (here, sentences). It is only because the successive words in sentences convey many possible meanings that we expect the internal vector that encodes them to span an increasingly larger portion of neural space.

Note that, while we predict increasing activity and intrinsic dimensionality due the recruitment of additional dimensions in normal sentences compared to Jabberwocky, there are a number of alternative hypotheses. Predictive coding theories of language processing (Shain et al., 2020; Heilbron et al., 2022; Goldstein et al., 2022) predict increased brain activations to surprising words, which in our setup should lead to normal sentences having the lowest responses. Likewise, if brain activity relates to processing difficulty (Just et al., 1996; Carpenter et al., 1999), then Jabberwocky should lead to the highest response. Thus, the predicted increase in dimensionality, greater for normal than for Jabberwocky sentences, should be found if and where brain signals are dominated by compositional semantics. Here, we do not make any specific claim about which characteristics of the stimuli influence the intrinsic dimensionality; future studies could test how parameters such as part-of-speech or conceptual specificity impact the dimensionality of neural representations.

### Multivariate decoding

We trained a logistic regression (Pedregosa et al., 2011) to separate normal and jabberwocky sentences at each time point using MEG and sEEG single-trial data. Such a decoding analysis informs us about whether and when our two conditions are differently represented in neural signals: if at time t the classifier reaches above-chance performance, it means that the brain (or the specific region of interest) segregates normal and Jabberwocky stimuli at this time. These classifiers were then tested at each other time point according to the temporal generalization method (King & Dehaene, 2014). This extension of the traditional within-time decoding analysis allows to test for the consistency of neural patterns over time: if a classifier trained at time t generalizes to time T, it means that the neural patterns is somewhat similar between time t and T. On the other hand, within-time decoding could be high at both t and T, but with no generalization between t and T. This would mean that the brain segregates normal and Jabberwocky stimuli at both time points, but with a different pattern of activations. In other words, the within-time decoding performance (trained and tested at the same time, i.e., the diagonal of the temporal generalization matrix) informs us about the content of brain signals, while the across-time decoding performance (trained and test at different times, i.e., the off-diagonal elements) tells us about the stability of these representations. Obviously, many different levels of internal representation may allow a classifier to categorize the incoming stimulus as a normal or a Jabberwocky sentence. However, the evolution of these representations over time should, at least partially, separate them.

Before training the classifiers, the data was subtracted from its median and scaled using the interquartile range, i.e. the range between the 1st quartile (25th quantile) and the 3rd quartile (75th quantile). We used a stratified k-fold crossvalidation procedure with 10 folds. We average the classifiers’ performances across all splits and report the average performance across subjects. We also stored the AUC for 100 permutations of the test labels in order to assess significance of the regional pattern regression analysis. To infer the overall tendency to generalize, we averaged each line of the temporal generalization matrix.

### Template regression

We trained a linear regression to predict empirical AUC matrices (averaged over patients) from templates. Specifically, each 601x601 matrix (we have 601 time points) was flattened to a 6012 vector, used in the regression analysis. For each template, we made a grid search to select the best parameters (delay and width, see Figure 7-1). We thus tested, for each empirical matrix, 100 candidates for each kind of template (phasic, ramping, sentence-final). The best template was the one with the highest likelihood in the regression model.

### Regional pattern analysis

Region of interests (ROI) were extracted from the Harvard- Oxford Cortical Atlas (Desikan et al., 2006). The whole decoding and pattern regression pipeline was repeated for each region. We only considered regions where at least three subjects had electrodes. The p values reported in this section have been FDR corrected at the region level.

### Neural Language Models

Neural networks have long been used to model natural language processing (Rumelhart & McClelland, 1986; Pater, 2019), and neural language models (NLMs) trained on next-word prediction have recently undergone a revival as models of human language processing (McClelland et al., 2020; Hale et al., 2021) and learning (Warstadt & Bowman, 2022); although see Lakretz, Desbordes, Hupkes, et al., 2021; Oh et al., 2022). Here we use NLMs as a testbed to check whether our hypotheses can be verified in a noise-free language processing system that has major implementational differences compared to biological neural networks. Several researchers have started to analyze activations from NLMs in an attempt to shed light on the neural codes for language (Tenney et al., 2019; Clark et al., 2019; Lakretz et al., 2019; Rogers et al., 2020), and the present work contributes to this field by introducing the temporal generalization method.

We report the results of home-trained character-based Transformers and Long-Short Term Memory (LSTM, (Hochreiter & Schmidhuber, 1997) models and of CamemBERT (Martin et al., 2020), a BERT (Devlin et al., 2019) model trained on a very large French dataset. The activations extracted from the CamemBERT models were obtained by giving the sentence piece by piece to the model and averaging the activation of each wordpiece composing a word. Thus, although the model is bidirectional we only gave it information about the past up to the current word.

The character-based LSTM models had 2 layers of 1024 units, while the character-based Transformer models had 12 layers and 768 units per layer. Both were trained on a 2GB sample of the French Wikipedia (Merity et al., 2016). We trained the models for 20 epochs, an initial learning rate of 20, a batch size of 128, a dropout rate of 0.2, and a sequence length of 35. At the end of an epoch, if no improvement was seen on the validation set, the learning rate was halved. For both LSTMs and Transformers models we used 10 instantiations of the model with different random seeds and report the performance averaged over all seeds. To obtain a single activation vector per word, we used the average activation evoked by each character belonging to the word for the character embedding layer, and the activations at the last character of the word for each upper layer. Untrained models were initialized with random weights (all sampled uniformly between -0.1 and 0.1), and directly underwent the same procedure for extracting their activations. We chose character-based models because word-based models cannot generalize to Jabberwocky (they only take trained words as input), whereas character-based models can take any string as input.

For the decoding analysis, we used a sample of 1000 sentences of each condition, generated using the same script as the subjects.

## Results

The present work aimed to address three main questions: how does the dimensionality of the neural representation evolve during sentence processing? Can phasic, ramping and sentence-final signals be disentangled in brain dynamics? Do they occur in separate brain regions? We start with a quick overview of the diversity of neural signals in our dataset. We then present the intrinsic dimensionality analysis, followed by multivariate decoding in artificial neural language models (NLMs) and real brain signals. Finally, we quantify the presence of each theoretical pattern in the empirical generalization matrices by means of multiple regression and replicate the decoding analysis in multiple brain regions. Overall, our results back the idea that learning language is associated with a predictable shaping of the representational manifold, such that meaningful sentences evoke ramping signals and a larger number of representational dimensions.

### Diversity of sEEG responses during sentence processing

Eleven patients with intracranial stereotactic electro-encephalography electrodes implanted for clinical purposes (sEEG, total n=2,243 electrodes; Figure 2-1) and simultaneously recorded magneto-encephalography (MEG, n=276 sensors per subject) read 240 sentences in rapid stream visual presentation (RSVP), with an SOA of 400 ms. Among them, half were normal French sentences, and the other half were the syntactically matched Jabberwocky sentences. Each sentence consists of eight words alternating between function words (determiners and auxiliary in position 1, 3, 5 and 7) and content words (nouns and verbs or their equivalent Jabberwocky pseudowords in position 2, 4, 6 and 8). These stimuli were mixed with 60 “string lists” of similar length, which consist of meaningless sequences of strings (Figure 2A) and were used as a low-level control. Specifically, as a localizer test, before all analyses, we selected language-selective channels using temporal cluster permutation test: only the channels with at least one significant cluster - after False Discovery Rate (FDR) correction at the electrode level - when comparing i) normal or Jabberwocky sentences to ii) string lists were kept for subsequent analyses.

We begin with a quick descriptive overview of the diversity of brain signals across electrodes and patients. Figure 3 illustrates some of these responses, both in the evoked broadband domain and in the high gamma frequency range (> 70 hertz). In each electrode, we assessed the variation of brain responses with our experimental conditions using a temporal cluster permutation test. Electrodes with at least one significant cluster - after False Discovery Rate (FDR) correction at the electrode level – when comparing i) normal and ii) Jabberwocky sentences, for either broadband or high gamma signals, were considered. We then manually selected representative electrodes from this pool. We do not claim these results to be exhaustive, rather we find it helpful to hold in mind these illustrative neural signals when examining the subsequent analyses. Unless specified otherwise, all subsequent statistical tests in this section are two-sample Wilcoxon-Mann-Whitney tests.

First, several visual electrodes exhibited a fast phasic broadband response, triggered indifferently by all visual stimuli, but modulated by stimulus length. As previously reported in other datasets (Agrawal et al., 2020; King et al., 2020; Woolnough et al., 2020), short function words triggered a smaller response than longer content words (p<0.01 based on the average activity between 50 ms and 300 ms following each stimulus presentation, electrode E6142, Figure 3-1A). Note that this channel is the only exception to the selection rule stated above: its activity did not differentiate between normal and Jabberwocky sentences, neither in broadband nor high gamma power.

Second, in the fusiform gyrus (FuG), the evoked responses of electrode E1042 (Figure 3A) was similar between normal and Jabberwocky, but significantly different from string lists (p<0.0001), consistent with this region’s sensitivity to written words and word-like stimuli (Woolnough et al., 2020). Interestingly, high gamma power from the same electrode exhibited phasic responses that differed for normal and jabberwocky (p<0.01, Figure 3A right), compatible with a role in lexical access (Woolnough et al., 2020).

Third, in regions such as the inferior parietal lobule (IPL, Figure 3B), high gamma responses to normal sentences showed a clear phasic effect. For example, in electrode E0064 it peaks 300ms after each content word. In this electrode, responses to Jabberwocky were very similar for the first pseudoword, but then quickly dropped.

While most electrodes exhibited stronger responses to normal sentences than Jabberwocky sentences, some responded specifically to Jabberwocky (e.g. superior temporal electrode E1263, Figure 3-1B). Such differences are compatible with the dual-route model of reading (Marshall & Newcombe, 1973; Jobard et al., 2003; Coltheart, 2005): while words evoke additional lexical, syntactic, semantic and, ultimately, compositional processes, pseudowords may also elicit specific processes associated for instance with attention and grapheme-phoneme conversion (Rumsey et al., 1997; Binder et al., 2003; Taylor et al., 2013).

Fourth, in inferior frontal gyrus (IFG, Figure 3C) and medial frontal gyrus (MFG, Figure 3-1C), we observed ramping responses: each additional content word led to an increase of the broadband and high gamma responses (linear regression on the difference in HG activity between normal and Jabberwocky for electrode E0878 (IFG) from 0 s to 3.5 s: slope=0.086, r=0.78, p<0.0001; and for electrodes E3652 (MFG) on the broadband: slope=4.99e-6, r=0.64, p<0.0001). This is compatible with a role in linguistic integration, in line with previous studies (Fedorenko et al., 2016; Nelson et al., 2017).

Last, in left superior frontal gyrus (SFG, Figure 3D) and right orbitofrontal cortex (OFC, Figure 3-1D), we observed sentence-final effects: for example, electrode E6062 (OFC), was mostly silent during the sentence, started to increase toward its ending, and exhibited a sharp peak more than 1 s after the last word’s onset (Figure 3-1D). On the other hand, electrode E0404 (SFG) responded similarly to both normal and Jabberwocky words, but these conditions ultimately diverged after the last word (Figure 3D).

Overall, the broad spectrum of functional responses illustrates the difficulty of interpreting the neural bases of language. Interestingly, a similar diversity of responses can be observed in individual units of deep language models (Figure 3-2). In the following sections, we use multivariate dimension reduction and decoding tools to evaluate whether our theoretical framework can account for the latent structure underlying these complex neural signals.

### Evaluating the intrinsic dimensionality hypothesis

As detailed in the introduction, our framework predicts that string lists, Jabberwocky sentences and normal sentences should lead to neural representations that systematically increase in their intrinsic dimensionality (ID). To compute the ID, we follow previous studies in neuroscience outside of the language domain (Gao et al., 2017; Sorscher et al., 2021; Elmoznino & Bonner, 2022) and use a method based on Principal Component Analysis (see Methods).

We performed this analysis on 400 ms time windows from -0.4 s to 5.2 s. Figure 4 shows the dimensionality estimate as a function of window onset. Three findings fit with our predictions. First, in all conditions for sEEG and for the normal text condition in MEG, the estimate increased with window onset, thus showing that as the successive words unfolded, the dimensionality of the brain signals increased (Pearson correlation with time, normal: R=0.98, Jabber: R=0.94; stringlist: R=0.89, p<0.0001 for each for broadband sEEG, Figure 4A; normal: R=0.91, p<0.0001, Jabber: R=0.50, p>0.05, stringlist: R=0.47, p>0.05 for MEG, Figure 4B, FDR corrected). Second, the intrinsic dimensionality was overall larger for normal sentences than for Jabberwocky sentences and string lists in the sEEG signals (p<0.001 for normal versus Jabber, p<0.001 for normal versus string lists, Wilcoxon-Mann-Whitney using the 4 s, FDR corrected, Figure 4A right). This was also the case for the MEG (p<0.001 for normal versus Jabber, p<0.001 for normal versus string lists, FDR corrected, Figure 4B right). Third, the difference increased as the sentence unfolded, as determined by a significant Pearson correlation between the difference (Normal – Jabber) and time (R=0.96, p<0.0001 for sEEG; R=0.92, p<0.0001 for MEG. We replicated these results with sEEG high gamma power and found highly similar results (Figure 4-1A).

**Figure 4:**
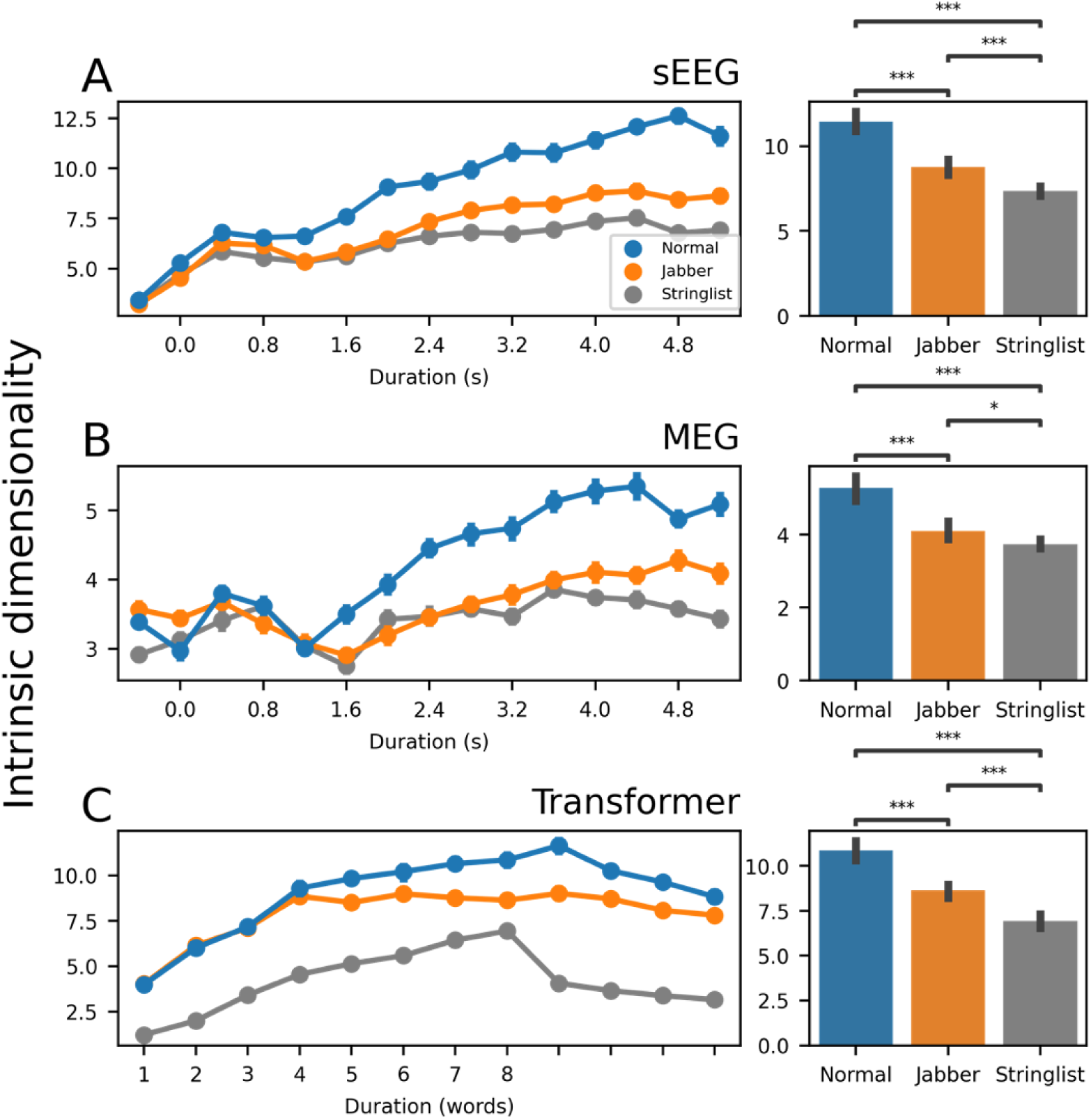
Intrinsic dimensionality is higher for normal sentences than Jabberwocky A. Intrinsic dimensionality computed from the broadband sEEG signals from all subjects as a function of the time window used (sliding time window of 0.4 s width). Right: Bar plot showing the intrinsic dimensionality computed using the whole sentence (a full 4 s time window). B, C, same analysis applied to MEG signals and Transformer activations (last layer, averaged over all stimuli for each condition). For the Transformer, the bar plot shows the intrinsic dimensionality computed using the 8 words time window. See Figure 4-1 for intrinsic dimensionality computed from high gamma sEEG, LSTM and CamemBERT See Figure 4-2 for the intrinsic dimensionality from untrained LSTM and Transformer models

Comparable effects were also observed in NLMs, such as causal Transformers (Figure 4C; p<0.001 for both pairwise comparisons; p<0.001 for all correlations), LSTMs (Figure 4-1B; p<0.001 for both pairwise comparisons; p<0.001 for all correlation) and CamemBERT (Figure 4-1C; p<0.05 for normal versus Jabber, p<0.001 for normal versus string list; p<0.0001 for all correlations). Crucially, untrained NLMs did not exhibit any significant differences between normal and Jabberwocky (Figure 4-2). Thus, in models, language learning is associated with a reshaping of the representational manifold, with an attribution of meaning to specific dimensions.

### Neural language models (NLMs) exhibit phasic, ramping and sentence-final responses

We next tested our second prediction, i.e. the existence of distinct phasic, ramping and sentence-final responses to sentences. To put these predictions to a test, we first evaluated whether these putative stages of semantic composition could be identified in NLMs using multivariate decoding and generalization. For this, we extracted the activations of each model in response to our stimuli and trained, at each word relative to sentence onset, a logistic regression across its artificial neurons to classify normal versus Jabberwocky sentences. We then evaluated these logistic regressions at each other time point, including the representations after the end of the sentence (obtained by feeding the model with four additional “space” tokens and extracting the corresponding activations). This temporal generalization analysis (King & Dehaene, 2014) resulted in a 12x12 training x testing matrix of classification scores summarizing i) where and when information distinguishing normal and Jabberwocky sentences is linearly represented in the network, and ii) whether the underlying representations change as the sentence unfolds.

The first computational step in NLMs is an “embedding” layer, where each vocabulary item (i.e., each character) is mapped onto a unique vector. Consequently, we expected temporal generalization to reveal a lexical signature in this embedding layer: i.e. a transient and phasic score rising after content words (Figure 2D). Our analysis confirmed this prediction: decoding led to a relatively small above-chance decoding performance at each content word (Figure 5A, mean AUC over all content words: 0.79, Wilcoxon-Mann-Whitney test against chance: p<0.0001). As expected, this signature disappeared for function words, as they were identical for normal and Jabberwocky sentences.

**Figure 5:**
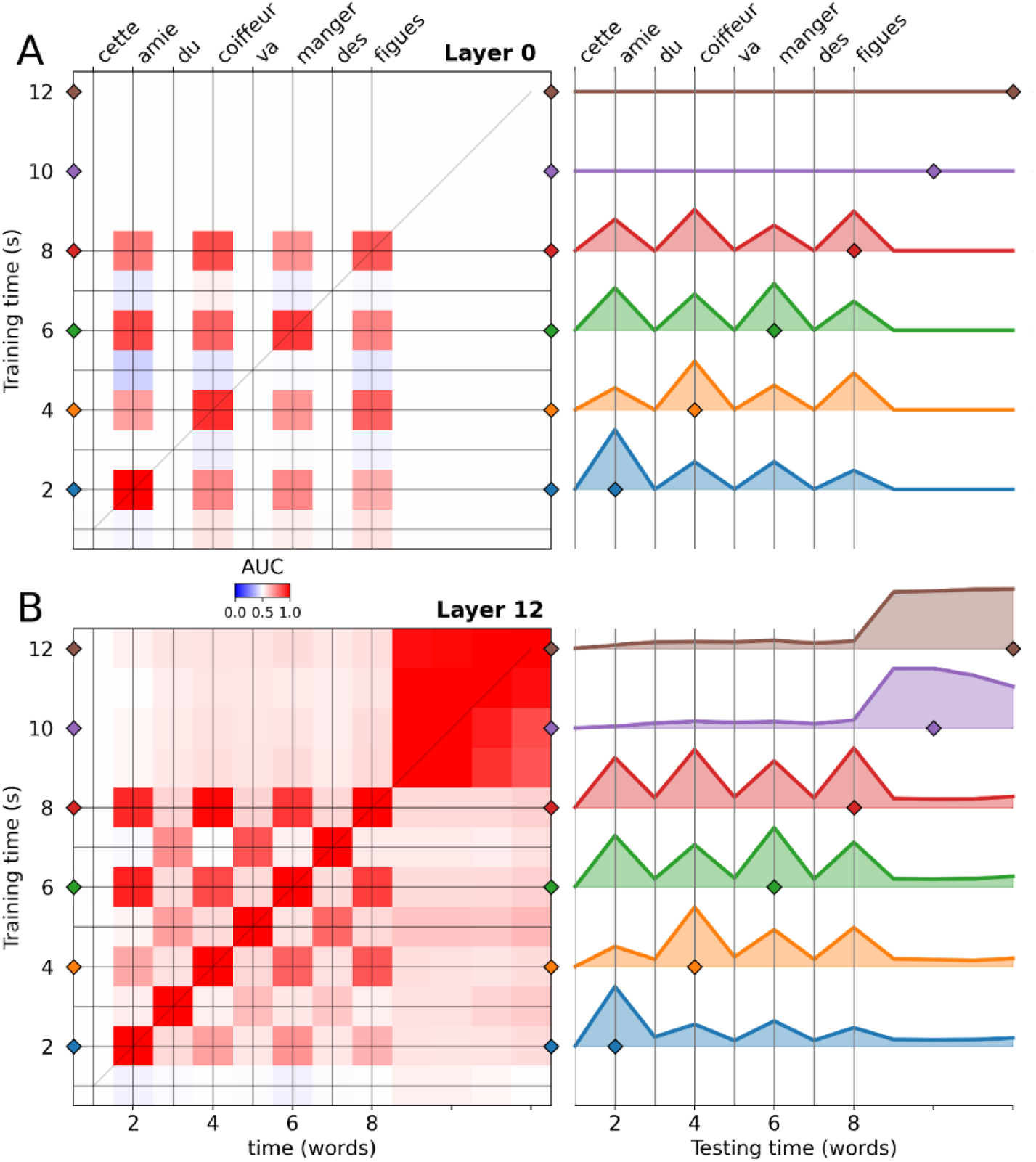
Decoding normal versus Jabberwocky sentences in neural language models shows lexical, persistent, and ramping patterns. Left: Temporal generalization matrices for a decoder trained to distinguish normal sentences from jabberwocky using the activity of the input word layer (top) and the final (12th) layer in a Transformer language model. The area under the curve (AUC) is the average over the 10 models trained on the same corpus but instantiated with different random seeds. Note that, in non-contextualized word embeddings, we only see the lexical pattern, whereas contextualized layers exhibit a superposition of multiple theoretical patterns. Although the performance on the diagonal is at ceiling (AUC=1), the generalization pattern is consistent with the ramping model. Right: Generalization of individual decoders, i.e., horizontal slices from the temporal generalization matrices on the left. These slices from each matrix show in more details the temporal dynamics of sentence processing. Filled lines show significant time points, tested with Wilcoxon-Mann-Whitney test against chance (0.5) and FDR correction. The first time point corresponds to the onset of the visual mask preceding the sentence. The onsets of successive words are marked by vertical grey lines. The small colored diamonds show the time where the decoders shown were trained. See Figure 5-1 for decoding normal versus Jabberwocky sentences in LSTMs and CamemBERT. See Figure 5-2 for the ramping tendency in LSTMs and Transformers NLMs.

The deep layers of NLMs integrate information from multiple tokens. Because of this integration of the preceding context, we expected temporal generalization to reveal an increasingly strong “square” of decoding performance. Indeed, we observed a temporal generalization not just between content words but also for function words as well as after the sentence. This was true for causal Transformers (Figure 5B), LSTMs (Figure 5-1A) and CamemBERT (Figure 5-1B). We show results for the last layer, where we found the overall decoding performance to be the strongest, nevertheless the earlier layers exhibited similar dynamics (Figure 5-2). Because the diagonal performance was at ceiling (AUC = 1) after the first content word, we could not directly test whether temporal generalization significantly increases over the course of the sentence, as predicted by a ramping processing stage (Figure 2D). However, the generalization performance of individual decoders (i.e. the average of each line from the matrix) increased over the course of the sentence, as revealed by a linear regression on the average of each line from word 1 to word 8 for layer 6, slope=0.017, r=0.78, p<0.0001). This finding suggests that NLMs demonstrate a ramping activity pattern that varies with semantic composition. Furthermore, this tendency to ramping increased in the upper layers of LSTM and causal Transformer models (Figure 5-2), suggesting that higher-level linguistic information is characterized by a stronger ramping signature.

Finally, looking at activations after the end of the sentence (i.e. when NLMs are input with spaces, upper right in Figure 5B), we observed a strong square generalization pattern (mean AUC for models trained and tested on tokens 9 to 12 : 0.97; Wilcoxon-Mann-Whitney test against chance: p<0.0001) that generalizes only modestly to the preceding words (mean AUC of classifiers trained on tokens 9 to 12, generalized to all preceding words: 0.59; Wilcoxon-Mann-Whitney test against chance: p<0.0001). Consistent with the predictions of a wrap-up processing stage, this result suggests that sentence-final representations partially differ from those generated during online sentence processing.

Overall, these analyses confirm that temporal generalization can isolate the three putative processing stages of semantic composition in NLMs: a phasic effect at the earliest processing stage of the network, and ramping and end-of-sentence effects at higher processing levels. In the next section, we apply these analyses on the sEEG and MEG responses to the same sentences, in order to test whether and where these processing stages occur in the human brain.

### Phasic, ramping and sentence-final patterns in time-resolved multivariate decoding

Applying multivariate decoding and temporal generalization to human sEEG (Figure 6A) and MEG recordings (Figure 6B) yielded a superposition of patterns. First, the sEEG and MEG diagonal decoding performance reached significance around 250 ms after word onset (Figure 6-1; cluster permutation test on sEEG: significant cluster from 0.69 s to 4.10 s, p=0.002, and MEG: first significant cluster from 0.63s to 1.10s, p=0.016), consistent with studies comparing ERPs evoked by words and pseudowords (Poldrack et al., 1999; Woolnough et al., 2020). Decoding reached a peak around 500 ms after stimulus onset, reaching up to 0.65 AUC +/- 0.03 (SEM) for sEEG (Wilcoxon-Mann-Whitney test against chance: p<0.01) and 0.58 AUC +/- 0.03 for MEG (Wilcoxon-Mann-Whitney test against chance: p<0.01). The multivariate pattern that separated normal and Jabberwocky was similar across the four positions where content words appear in the sentence, resulting in a 4-by-4 grid of decoding generalization, similar to the one observed in neural language models (Figure 5). The resulting grid pattern means that even though the neural activity evolved over the course of the sentence, after each content word it transiently reached a similar state that dissociated normal and Jabberwocky sentences. This phasic pattern is consistent with a lexical process (Figure 2D).

**Figure 6:**
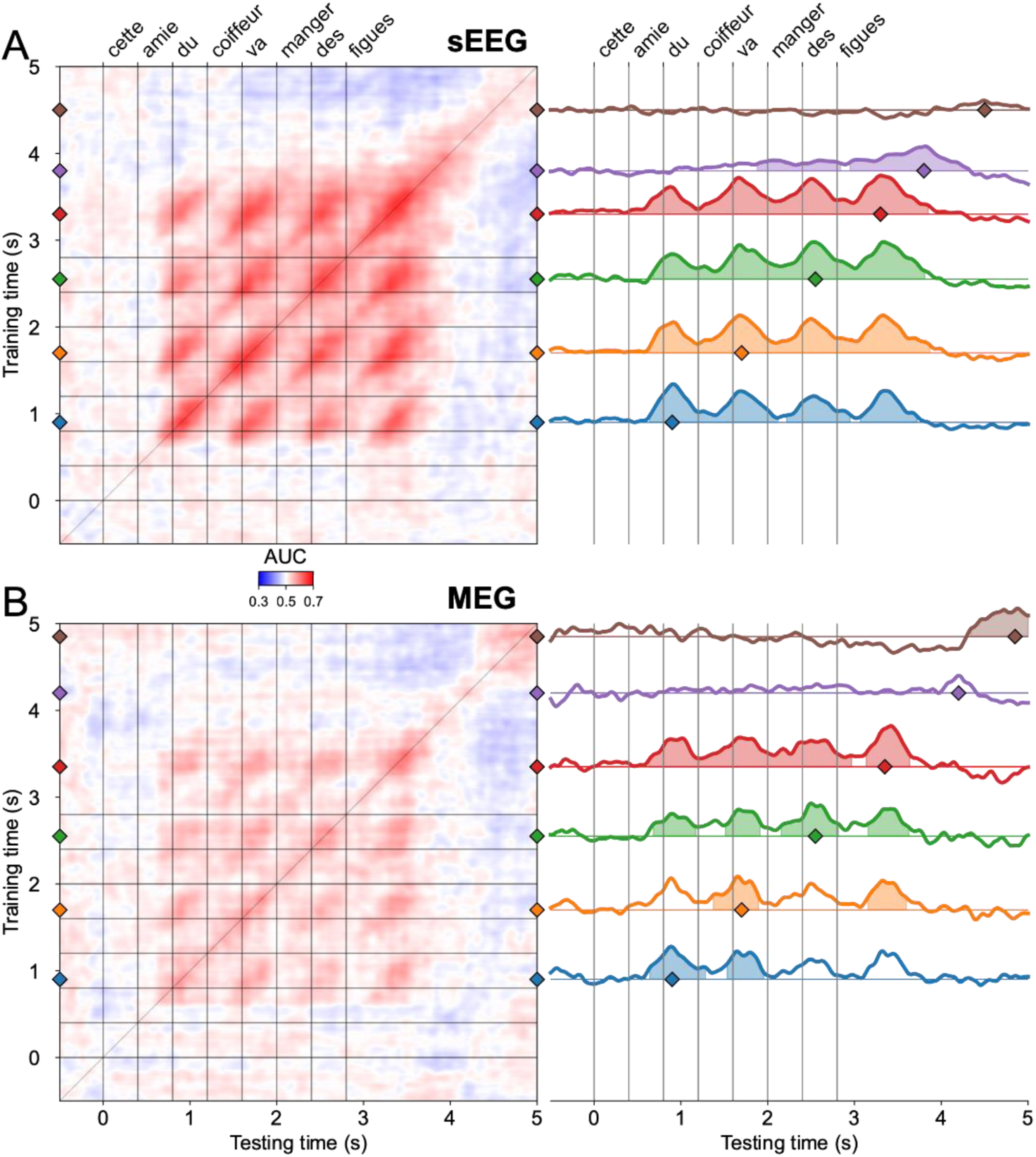
Decoding normal from Jabberwocky in human sEEG and MEG shows phasic, ramping and sentence-final patterns Left: Temporal generalization matrices for a decoder trained to distinguish normal sentences from jabberwocky using in human sEEG (A) and MEG (B). The AUC is the average over the 11 subjects for sEEG and 9 subjects for the MEG. Right: Generalization of individual decoders, i.e., horizontal slices from the temporal generalization matrices on the left. These slices from each matrix show in more details the temporal dynamics of sentence processing. Filled lines show significant time points, tested with cluster permutation test and FDR correction. The first time point is the onset of the visual mask preceding the sentence. Each vertical grey line is a word onset. The small colored diamonds show the time where the decoders are trained. See Figure 6-1 for the diagonal decoding performance for normal versus Jabberwocky sentences in human sEEG and MEG

Second, to examine the presence of ramping effects, we tested whether decoding and generalization performance (average performance of a decoder over all timepoints) increased with sentence unfolding (from 0.4 s and 4 s). We found a positive effect in sEEG diagonal performance (linear regression on the average AUC across subjects: slope=0.013, r=0.39, p<0.0001), and the generalization performance (slope=0.0068, r=0.61, p<0.0001). In MEG, there was a significant effect for the diagonal performance (slope=0.0053, r=0.25, p<0.0001), but no effect for the generalization performance (slope=0.000072, r=0.014, p=0.81).

Finally, decoding performance remained significant for more than one second after the end of the sentence (cluster permutation test on sEEG: significant cluster from 0.69 s to 4.10 s, p<0.01, the last word’s onset being at 2.8 s, Figure 6-1). For example, the purple line in Figure 6A corresponds to a sEEG classifier trained 1 s after the last word’s onset. Despite being trained this late, it reached an AUC of 0.59+/-0.03 and generalized to a few seconds before, with a clear ramping pattern (cluster permutation test: 2 significant clusters from 1.9 s to 2.8 s, p=0.014, and from 3 s to 4.3s, p<0.01). Similarly, the MEG classifier trained 2 s after the onset of the last word (t=4.9 s, Figure 6B brown line), hence much later than sensory and lexical processes, still showed a high decoding performance (training time AUC =0.56+/-0.03, cluster permutation test: one significant cluster from 4.3 to 5 s, p<0.01), now quite restricted over time and hence supporting the sentence-final wrap-up hypothesis (Figure 2D).

To summarize, at the whole brain level, we observed a linear superposition of the 3 patterns (Figure 2D) hypothesized to participate in semantic composition. Together, these results suggest that the brain integrates semantic information across multiple words and, after the end of the sentence, reaches a state that still differentiates normal and Jabberwocky sentences for a long period of time.

### Superposition and regional specialization of phasic, ramping and sentence-final effects

To quantify the extent to which each of the three dynamic patterns was present in the empirical generalization matrices, we fit a linear regression using the (linearized) template matrices as predictors:

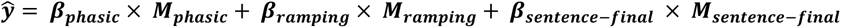

In this equation, ŷ is the predicted AUC matrix (average of all subjects), *M_phasic_*, *M_ramping_* and *M_sentence-final_* are the template matrices shown in Figure 2D, and the betas are the corresponding estimated coefficients. To account for varying time delays and intrinsic dynamics, we performed a systematic grid search over template matrices, individually varying the onset and width of each peak (Figure 7-1). The model with the highest likelihood was selected. We thus obtained a beta coefficient for each template, quantifying the degree to which the dynamic pattern was present in the empirical matrix. Note that this analysis is coarse: because we average patients’ data before fitting the regression, only consistent patterns across patients will show up. The reason for this averaging is the small number of patients (11) and the fact that each patient does not have electrodes in every region, thus making the total number of data points too small for fitting a regression per patient followed by statistical testing across patients. Instead, we assess statistical significance with permutation tests (by shuffling the classifier’s labels at the time of training the decoder).

**Figure 7:**
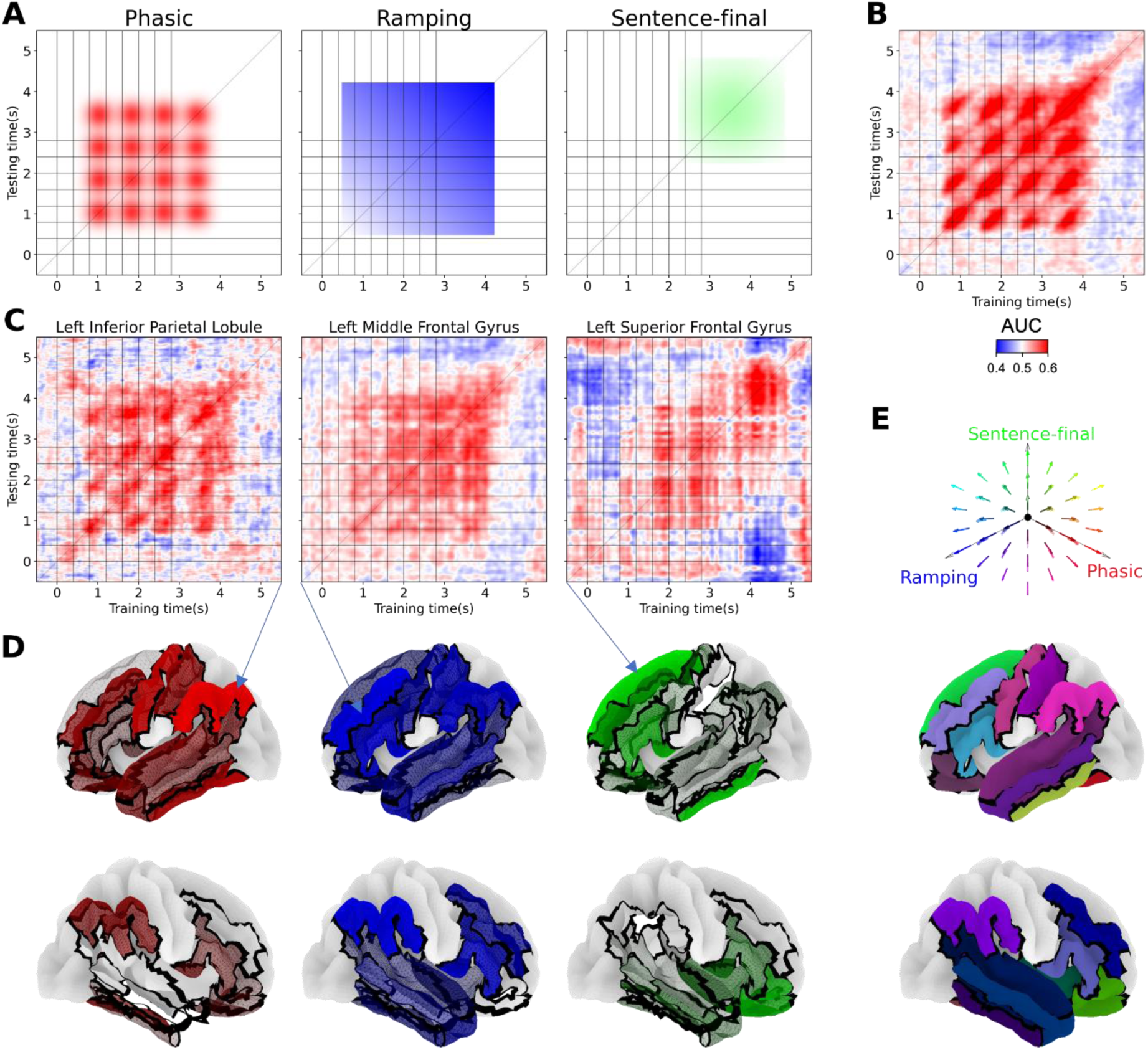
Phasic, ramping and sentence-final patterns are found in to varying degrees in each region in human sEEG A. Template matrices selected by the grid search for whole-brain human sEEG. For each template, 20 different template matrices with varying delays and widths (Figure 7-1) were tested against the data, and the best fit was kept. B. Empirical matrix for whole-brain human sEEG. C. Empirical matrices in human sEEG for three most relevant ROIs. Arrows show the corresponding brain regions. D. Surface brains maps showing the strength of the temporal generalization patterns for phasic (red, left), ramping (blue, middle), and sentence-final (green, right) processes in each ROI. The brain maps show the corresponding red, blue or green value, with a transparency value proportional to the regression coefficient of the corresponding pattern. E. Surface brain map showing the combined strength of the 3 patterns in each ROI. The color of each ROI reflects the red, blue, and green values of the phasic, ramping and sentence-final patterns. See Figure 7-1 for illustrative templates used in the grid search for the template regression analysis. See Figure 7-2 for the lack of syntactic modulation of the ramping pattern.

Applying this method to the whole brain temporal generalization matrix (Figure 6B), we obtained significant coefficients (i.e., bigger than the coefficients fit on the AUC matrices from shuffled labels) for the three patterns: *β_phasic_* = 0.11 (p < 0.01), *β_ramping_* = 0.06 (p < 0.01), *β_sentence-final_* = 0.12 (p < 0.01), confirming the findings of the previous section. The corresponding template matrices are shown in Figure 7A.

To evaluate whether the three dynamics revealed by temporal generalization arise from distinct brain regions, we repeated this regression analysis on subsets of electrodes belonging to anatomically defined regions of interest (ROIs), and predicted to be involved in distinct language-specific processes (Hickok & Poeppel, 2007; Friederici, 2011; Hagoort, 2019; Matchin & Hickok, 2020). We then plotted these results for each ROI by assigning a red, green and blue value corresponding respectively to the beta (normalized across regions to be between zero and one) of the phasic, ramping and sentence-final patterns. Example empirical matrices are shown in Figure 7C and the resulting whole brain maps in Figure 7D and 7E.

First, we expected the ventral occipito-temporal visual pathway, and particularly the FuG, to exhibit phasic (lexical) effects, with no ramping or delayed patterns. Our results are broadly consistent with this prediction as shown by the strong presence of the lexical component (*β_phasic_* = 0.84, p < 0.01), but not ramping (*β_ramping_* = 0.14, p < 0.12) nor sentence-final (*β_sentense-final_* = 0.07, p = 0.09) components in the FuG (Figure 7D and 7E). To verify that the ramping pattern was not present in these regions, we checked that the performance increased only marginally and non-significantly over the course of the sentence (linear regression AUC for each subject: average slope=0.0018, Wilcoxon Mann-Whitney test against null slope: p=0.43 for the left precentral gyrus).

Second, we expected the MTG and STG, to be responsible for accessing lexical representations (Hart et al., 2000; Binder et al., 2003; Tranel, 2009) and starting to compose them according to sentential context (Lau et al., 2008; Pallier et al., 2011; Price et al., 2015, 2016), thus showing a combination of the phasic and ramping patterns. Our results are consistent with this prediction (left STG: *β_phasic_* = 0. 54, p < 0. 01, *β_ramping_* = 0. 53, p < 0. 01, *β_sentence-final_* = 0.17, p = 0.1; left MTG: *β_phasic_* = 0.42, p < 0. 01, *β_ramping = 0.6_*, p < 0. 01, *β_sentence-final_* = 0.12, p = 0.08): the normal versus jabberwocky decoding results revealed a strong response starting 200 ms and peaking 400 ms after each content word onset (e.g. peak performance for the first word at t=0.8 s: mean AUC=0.56 +/- 0.02 Wilcoxon-Mann-Whitney test against chance: p<0.01), with good generalization to all words in the sentence along with an increase of performance over time (linear regression for each subject: average slope=0.0039, Wilcoxon Mann-Whitney test against null slope: p<0.01) and an increase of the generalization performance (average slope=0.0047, Wilcoxon Mann-Whitney test against null slope: p<0.01). Interestingly, parietal regions such as the IPL showed a similar pattern (figure 7C left): *β_phasic_* = 1, p < 0. 01, β_ramping = 0.83_, p < 0. 01, *β_sentence-final_* = 0.18, p = 0.07, confirming their involvement in word composition (Bemis & Pylkkänen, 2013; Price et al., 2015, 2016).

Third, we expected the prefrontal cortex to be more specifically involved in combinatorial computations (Hagoort, 2005; Pallier et al., 2011; Friederici, 2011; Fedorenko et al., 2016; Nelson et al., 2017; Matchin & Hickok, 2020). In the left IFG (*β_phasic_* = 0.2, p = 0.08, *β_ramping_* = 0.85, p < 0. 01, *β_sentence-final_* = 0.69, p < 0. 01) and in the left MFG (Figure 7C middle; *β_phasic_* = 0.64, p < 0. 01; *β_ramping_* = 1, p < 0. 01, *β_sentence-final_* = 0.58, p < 0. 01), we observed a ramping activity profile, starting around 350 ms after the first content word’s onset, and increasing without discontinuity until 4.2s, i.e., 1.4 s after the last word onset (linear regression for each subject: average slope=0.0065, Wilcoxon Mann-Whitney test against null slope: p<0.001 for left IFG). The ramping pattern was also visible in the generalization performance (average slope=0.0054, Wilcoxon Mann-Whitney test against null slope: p<0.01). Similar ramping profiles were found in the right IFG and MFG (Figure 7D and 7E), but the evidence for phasic and sentence-final patterns was weaker. This activity profile is consistent with a linear integrator (Pallier et al., 2011; Fedorenko et al., 2016), whereby I/MFG would combine each incoming word with the previous ones.

Last, in the right OFC, (*β_phasic_* = 0.26, p=0.15, *β_ramping_* = 0., p = 0.96, *β_sentence-final_* = 0.71 p < 0.01), and, the left SFG (Figure 7C, *β_phasic_* = 0.09, p = 0.47, *β_ramping_* = 0.34, p = 0.05, *β_sentence-final_* = 1, p < 0.01), decoding performance stayed at chance level for the most part of the sentence, but significantly increased after the last word, and stayed above chance until 1.6 s after the end of the sentence (cluster permutation test on the diagonal performance: single significant cluster from 3.9 s to 4.4 s, p<0.01 for left SFG and single significant cluster from 4.0 s to 4.4 s, p<0.01 for right OFC). This sentence-final effect is consistent with a wrap-up process.

In sum, we successfully identified a set of regions exhibiting signatures of phasic, ramping and sentence-final processes, and thus provide a path to a systematic decomposition of sentence composition in the brain.

## Discussion

To clarify how sentences are composed by the human brain, we introduced and tested a simple yet powerful vector coding framework in NLMs and human electrophysiological recordings. First, we predicted that the dimensionality of brain signals should increase as successive words get added to an evolving representation of sentential meaning, and that this effect should be larger for meaningful than for meaningless materials (Jabberwocky or lists of meaningless strings). Indeed, we found that representations of meaningful sentences evoke neural signals of higher dimensionality than Jabberwocky sentences or string lists. Noteworthy, there was an increase of dimensionality with time in all conditions but, crucially, it was highest for normal sentences. These effects were absent in untrained NLMs, supporting the idea that, with learning, coding dimensions are assigned distinctive meaning in distributed semantic spaces.

To further characterize the dynamics of brain activity during sentence processing we used multivariate decoding and temporal generalization, a method that has recently been advocated for in the context of language processing (Fyshe, 2020; He et al., 2022). The results indicate that the representations generated in brains and NLMs follow similar dynamics, despite their differences in implementation and timescale. Specifically, we observed three distinct dynamic signatures: phasic, ramping and sentence-final, which we interpret as the reflection of single-word processing, multi-word composition, and sentence wrap-up, respectively. ROI analysis showed that the FuG, exhibited pure lexical patterns, confirming its role in written word identification. On the other hand, regions such as MTG, STG and IPL, displayed mixed lexical and ramping patterns, suggesting that access single-word meanings and start to combine them. Frontal regions had mixed signatures of ramping and sentence-final (for IFG), as well as a phasic component (for MFG), or a pure sentence-final effect (for right OFC and left SFG). Overall, we observe a wide variety of combinations of each signature, witnessing the dynamic flow of information during sentence processing. Although this pattern-based analysis is coarse and is correlational rather than causal, it is corroborated by single-channel evoked activities (e.g. Figure 3) and fits with the previous literature.

Lexical access, in particular, has been studied extensively and is thought to be supported by the FuG and temporal regions, where we found strong phasic signatures. Curiously, this was also the case of parietal regions such as IPL and the pre and postcentral gyri, suggesting a stronger involvement in lexical access than previously thought. The ramping pattern also appeared as a marker of compositional processes in several previous studies. Pallier and colleagues (2011) observed that fMRI activity in IFG and posterior Superior Temporal Sulcus (STS) increased in direct proportion to the number of elements in the current syntactic phrase, for both normal text and Jabberwocky. They proposed a simple model in which each consecutive word or phrase adds a fixed amount of activity to a compositional representation which therefore builds up across time. Nelson and colleagues (2017) and Fedorenko et al. (2016) then showed, with the higher resolution of intracranial EEG, that high-gamma activity does indeed increase after each word in a constituent word phrase.

The exact computational role of this ramping activity is, however, still unknown. Due to the engagement of additional semantic processes, the neural assemblies recruited during the processing of normal sentences should be larger and the neural dynamics richer, compared to Jabberwocky. Computational models of the neural encoding of compositional structures (Smolensky, 1990; Plate, 1995; Gayler, 2004) indeed predict that the neural code can be characterized as a sum of representations for each constituent (technically, a sum of tensor products of the vectors representing each word’s role and filler) and should therefore increase as their number increases. Nevertheless, direct evidence for such a neural code is still missing. Testing whether brain activity during sentence processing contains signatures of tensor product representations is a promising avenue for future work.

The sentence-final pattern seen in SFG and OFC may be associated with several cognitive processes, collectively dubbed wrap-up processes. The classical view holds that sentence wrap-up effects reflect higher-level integration and if necessary, reanalysis and conflict resolution of the multiple possible meanings of words (Just & Carpenter, 1980; Molinaro et al., 2008). Until recently, few neuroimaging studies examined sentence-final activations, for fear that wrap-up effects would confound regular processes happening at the last word (Stowe et al., 2018). The present study shows one way that they can be disentangled.

Note that, here, we also used varied syntactic structures (Figure 2C) as an initial attempt to look for modulations of the dynamic patterns by syntax. However, no such modulations were found (Figure 7-2), suggesting that the integrative processes are similar in the three structures used. This does not preclude a modulation in more complex structures, for example sentences including adverbial phrases or embedded clauses.

The present intrinsic dimensionality hypothesis stems from much research in the past decades, which has emphasized how biological neural networks use high-dimensional vector spaces to encode complex structures (Quiroga et al., 2005; Tyukin et al., 2019; Gorban et al., 2019; Calvo Tapia et al., 2020). For instance, human MEG signals can be decomposed into several dimensions that reflect the various contents of a visual sequence (Liu et al., 2019; Quentin et al., 2019; Al Roumi et al., 2021). Similarly, recordings of thousands of monkey prefrontal neurons can be decomposed into orthogonal vector subspaces storing the successive elements of a spatial sequence in working memory (Xie et al., 2022). The present intrinsic dimensionality analysis extended this idea to sentences. Non-linear alternative measures of dimensionality are available (Granata & Carnevale, 2016; Facco et al., 2017; Landa et al., 2021), but the advantages of the measure of intrinsic dimensionality used here are its simplicity and wide acceptance.

Distributed word representations from NLMs were shown to align with cortical responses to words by means of linear encoding models (Huth et al., 2016; Jain & Huth, 2018; Toneva & Wehbe, 2019; Caucheteux & King, 2020; Caucheteux et al., 2021; Schrimpf et al., 2021; Goldstein et al., 2022). Such partial similarities in brain activity, in performance on various linguistic tasks (Otter et al., 2021), and in error types (Coenen et al., 2019; Goldberg, 2019; Jawahar et al., 2019; Lakretz et al., 2020; Lakretz, Desbordes, King, et al., 2021) provide an interesting, although incomplete, means to study neural representations of sentences during language processing in a system which is (1) less noisy compared to brain data, (2) fully accessible to manipulation and recordings, and (3) not a priori tied to a normative linguistic theory but rather, learned from large-scale linguistic corpora. The present decoding approach shows that the similarities between brains and artificial neural networks lie not only in their representations, but also in their dynamics (i.e., the order in which the representations are combined with one another). Perhaps even more surprisingly, the resemblance holds for non-canonical stimuli such as Jabberwocky sentences, which are out-of-domain for both brains and NLMs. Of note, the interpretation of the sentence-final pattern in the models is not straightforward and we remain careful to interpret it as a wrap-up process.

A limiting factor in our work is the limited coverage of some brain regions (e.g., STS). Furthermore, we had to pool together electrodes in relatively large brain regions in order to achieve decent decoding performance. This choice greatly limited the spatial accuracy of our analyses. For example, subregions of the IFG are known to have functional specializations and inter-individual variability (Fedorenko & Blank, 2020): e.g., the pars triangularis (BA45) may be most sensitive to syntax (Nelson et al., 2017). Similarly, we had to aggregate the temporal pole, which has been associated with 2-words composition (Bemis & Pylkkänen, 2011; Fyshe et al., 2019; Pylkkänen, 2019), with the temporal gyri. Higher-resolution recordings, for instance using smaller electrode arrays (Szostak et al., 2017; Steinmetz et al., 2021), will be needed in order to further study the functional specialization of these regions. Furthermore, our study considers semantic composition in the broadest sense and does not afford any claim regarding the specific subprocesses underlying the observed dynamics.

Taken together, our results suggest that a succession of processing stages, separated by their distinct brain signatures, underlie the composition of sentence-level semantics. They allow us to speculate that incoming lexical information arising from the FuG (Woolnough et al., 2020) is first passed to the temporal lobe and IPL, where semantic information is accessed, stored, and begins to be combined. The I/MFG exhibits relatively selective ramping and sentence-final signals, suggesting that it may play a key role in merging individual words into constituents that encode their compositional meaning. The final sentential representation may then stay present in the activity pattern of the left SFG and right OFC for several seconds – a duration which might have been extended if we had presented multiple sentences forming part of a larger discourse. Finally, the construction of these compositional meanings is associated with an increased dimensionality of the representations. These results bring us one step closer to understanding how the human brain composes and understands sentences.

## Acknowledgements

Researchers from the Institute of Language, Communication and the Brain (ILCB) were supported by grants ANR-16-CONV-0002 and ANR-11-IDEX-0001-02 (A*MIDEX). Christian Bénar was partly funded by a FLAG ERA/HBP grant from Agence Nationale de la recherche “SCALES” ANR-17- HBPR-0005. Stanislas Dehaene was funded by INSERM, CEA, Collège de France, the Bettencourt-Schueller foundation and an ERC grant “NeuroSyntax”. Data acquisition was performed on a platform member of France Life Imaging network (grant ANR-11-INBS-0006).

---

**Figure 2-1:**
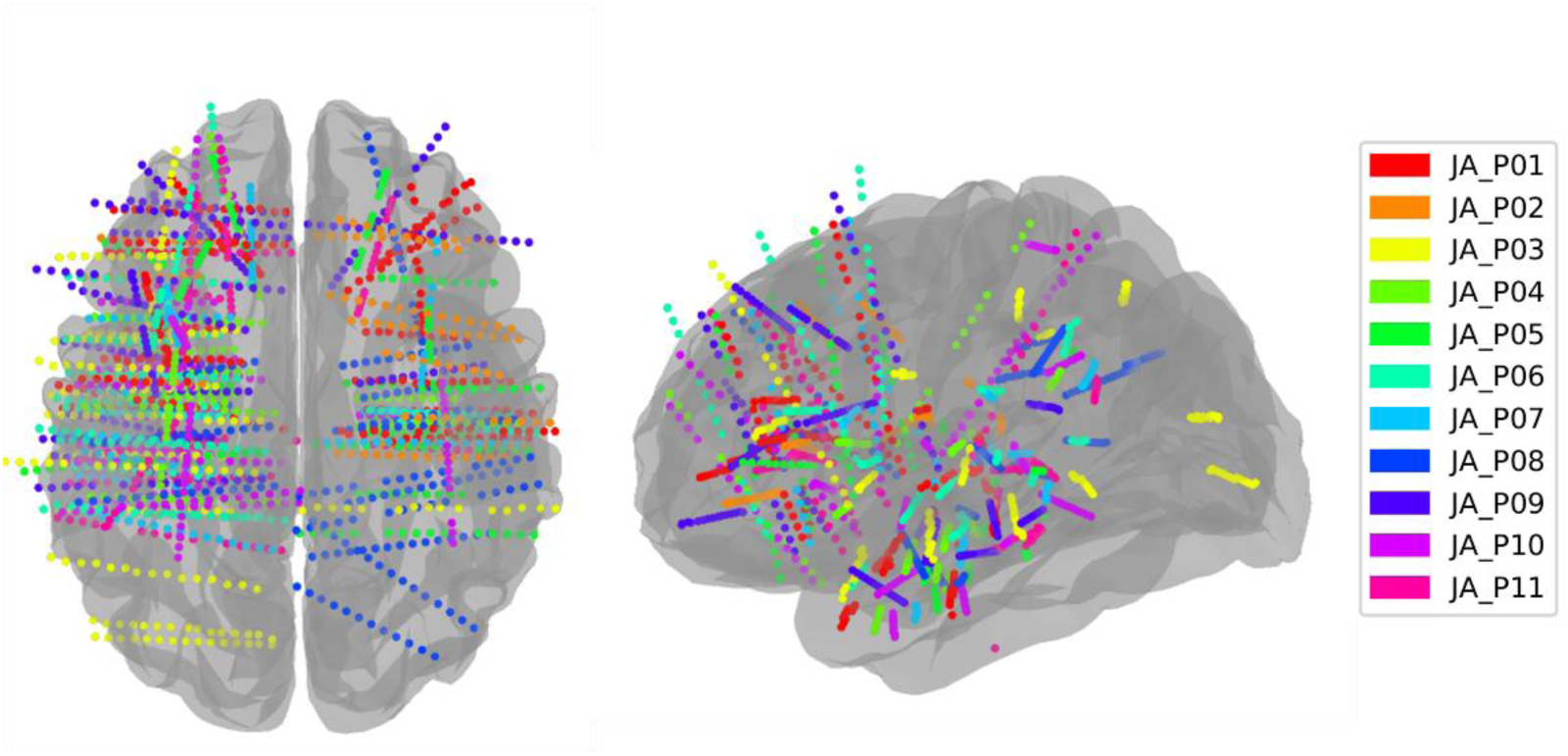
Intracranial electrode coverage. Electrode location in Montreal Neurological Institute (MNI) space. Each patient is shown with a different color.

**Figure 3-1:**
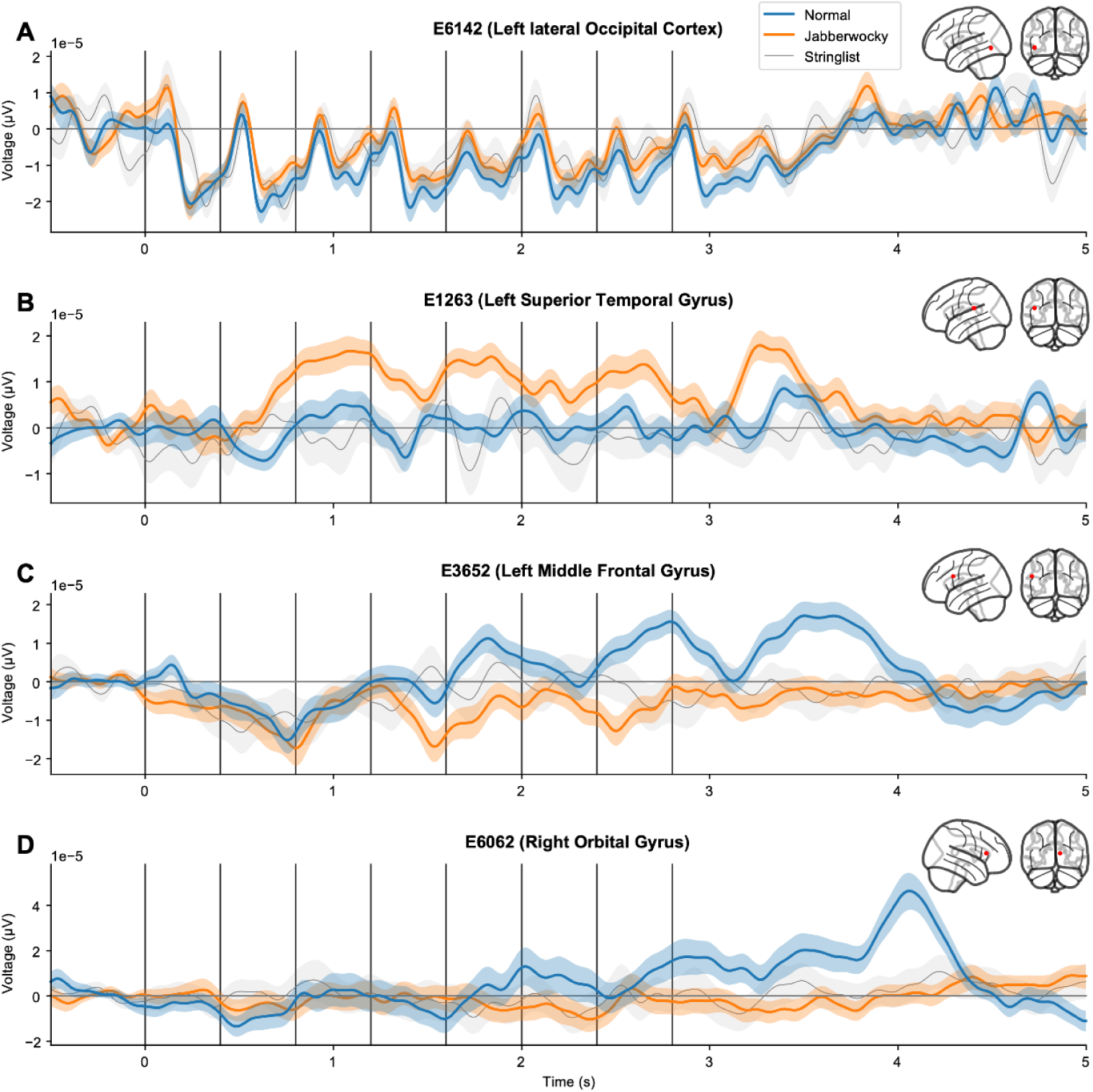
Additional illustrative profiles of human sEEG responses. Four examples of electrodes responding to normal sentences (blue), Jabberwocky sentences (orange), and string of consonants (gray). Each line shows the local field potential (Voltage) for each electrode.

**Figure 3-2:**
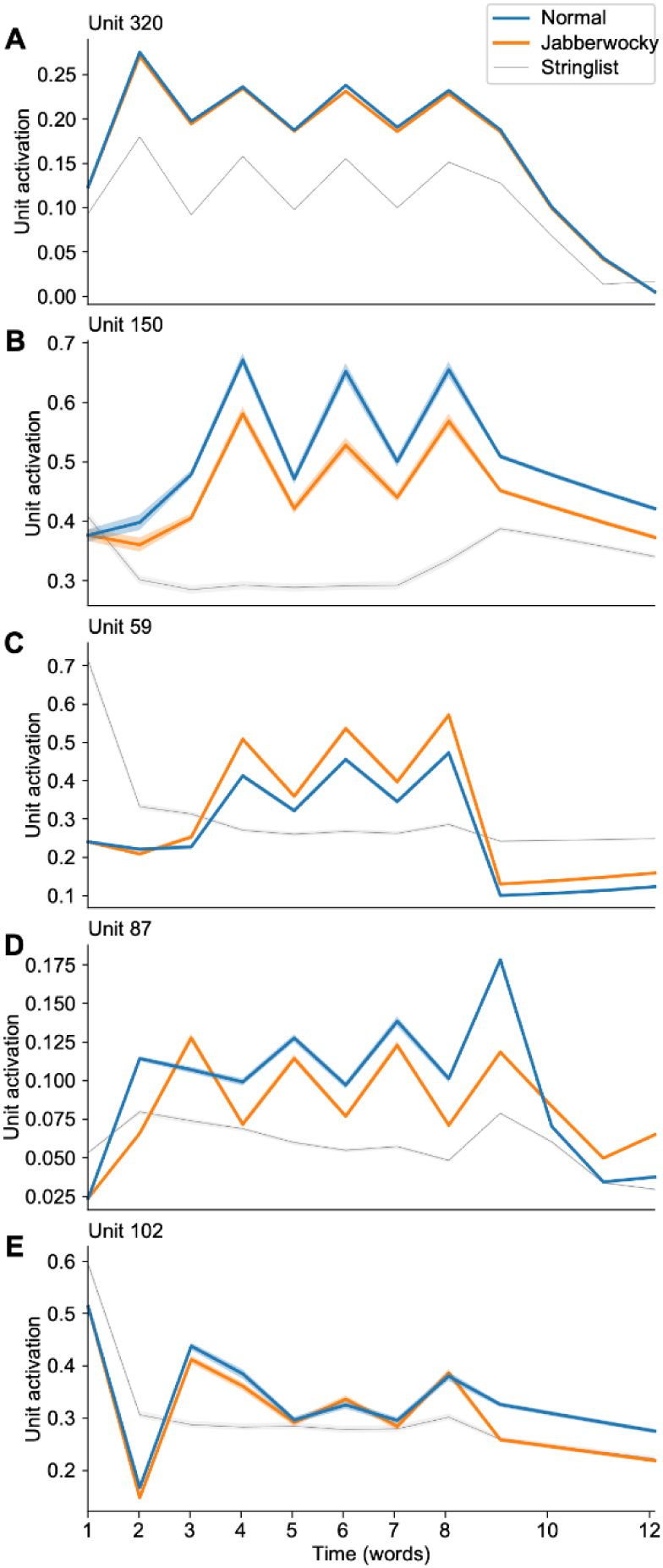
Average activation from hand-picked units in the Transformer’s 12th layer, showing interesting dynamics, sometimes surprisingly similar to human sEEG activations. A. Orthographic effect. Word length affects the activations (higher for longer words). B. Lexical effect. Activation differs between normal and Jabberwocky after the first content word. C. Same as B but with higher activations in the Jabberwocky condition. D. Ramping effect. The difference between normal and Jabberwocky increases over the course of the sentence. E. Sentence-final effect. Although the activations are similar during the sentence, normal and Jabberwocky diverge after the presentation of the last (8th) word.

**Figure 4-1:**
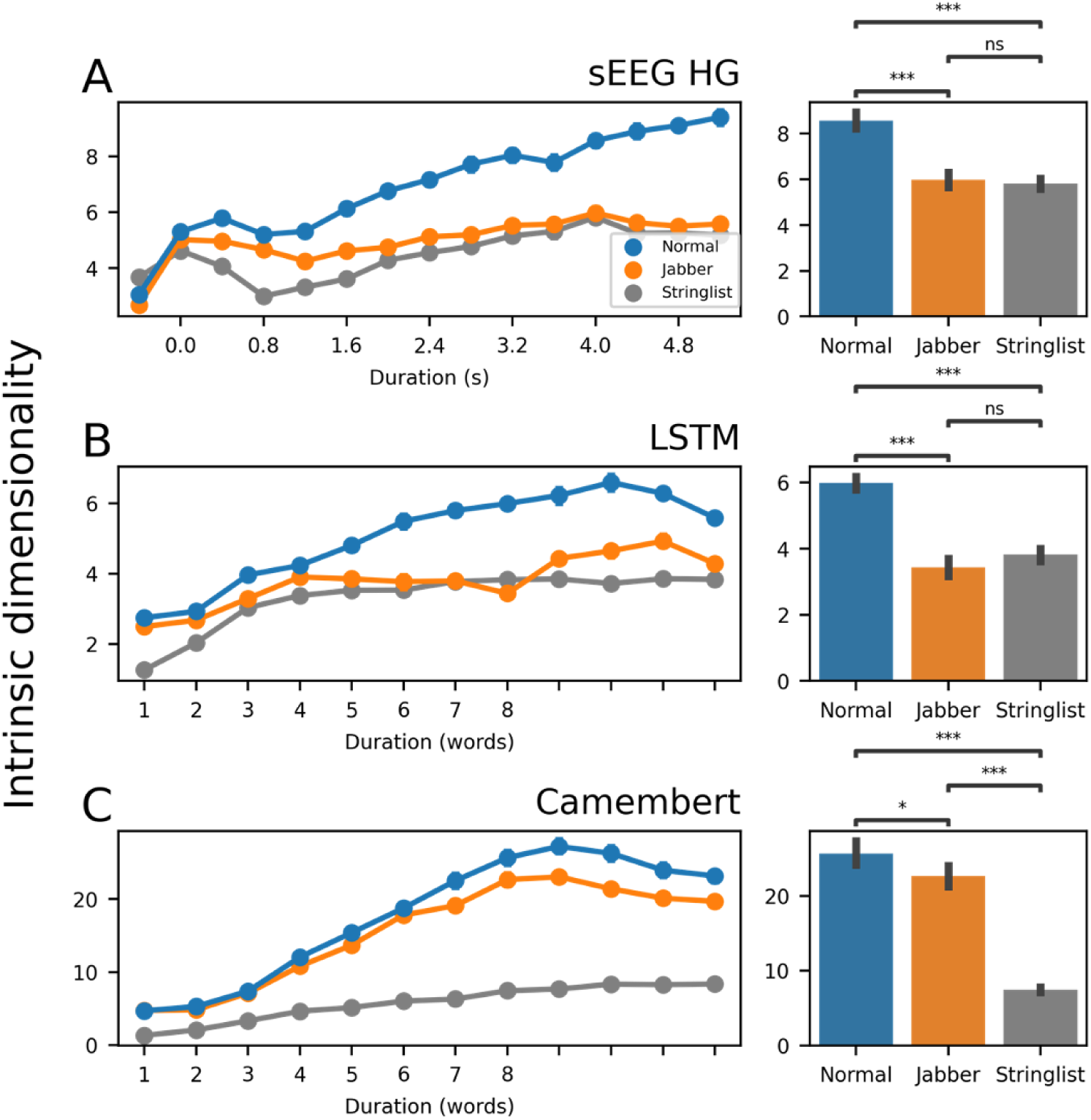
Intrinsic dimensionality is higher for normal sentences than Jabberwocky A. Intrinsic dimensionality of the sEEG high gamma signals from all subjects, as a function of the time window used (sliding time window of 0.4 s width). Right: Bar plot showing the intrinsic dimensionality computed using the whole sentence (4 s time window). B, C, same analysis applied to LSTM and CamemBERT activations (last layer, averaged over all stimuli for each condition). The bar plots show the intrinsic dimensionality computed using the 8 words time window.

**Figure 4-2:**
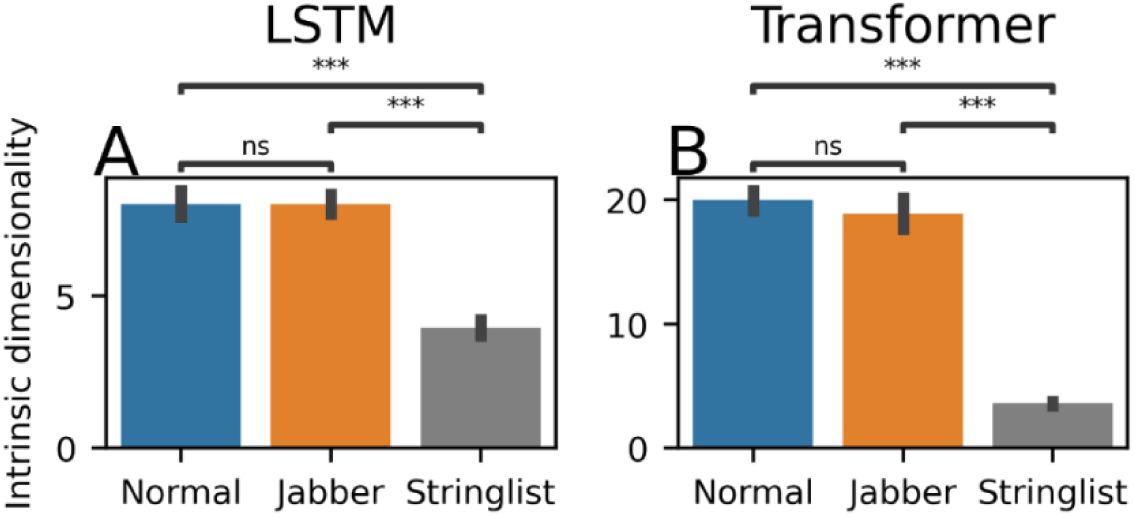
Untrained language models do not exhibit a larger intrinsic dimensionality for normal versus Jabberwocky sentences. The figure shows the Intrinsic dimensionality computed using the whole sentence (8 words) time window for an untrained LSTM (A) and an untrained Transformer (B) e was no significant difference between the normal and Jabberwocky conditions.

**Figure 5-1:**
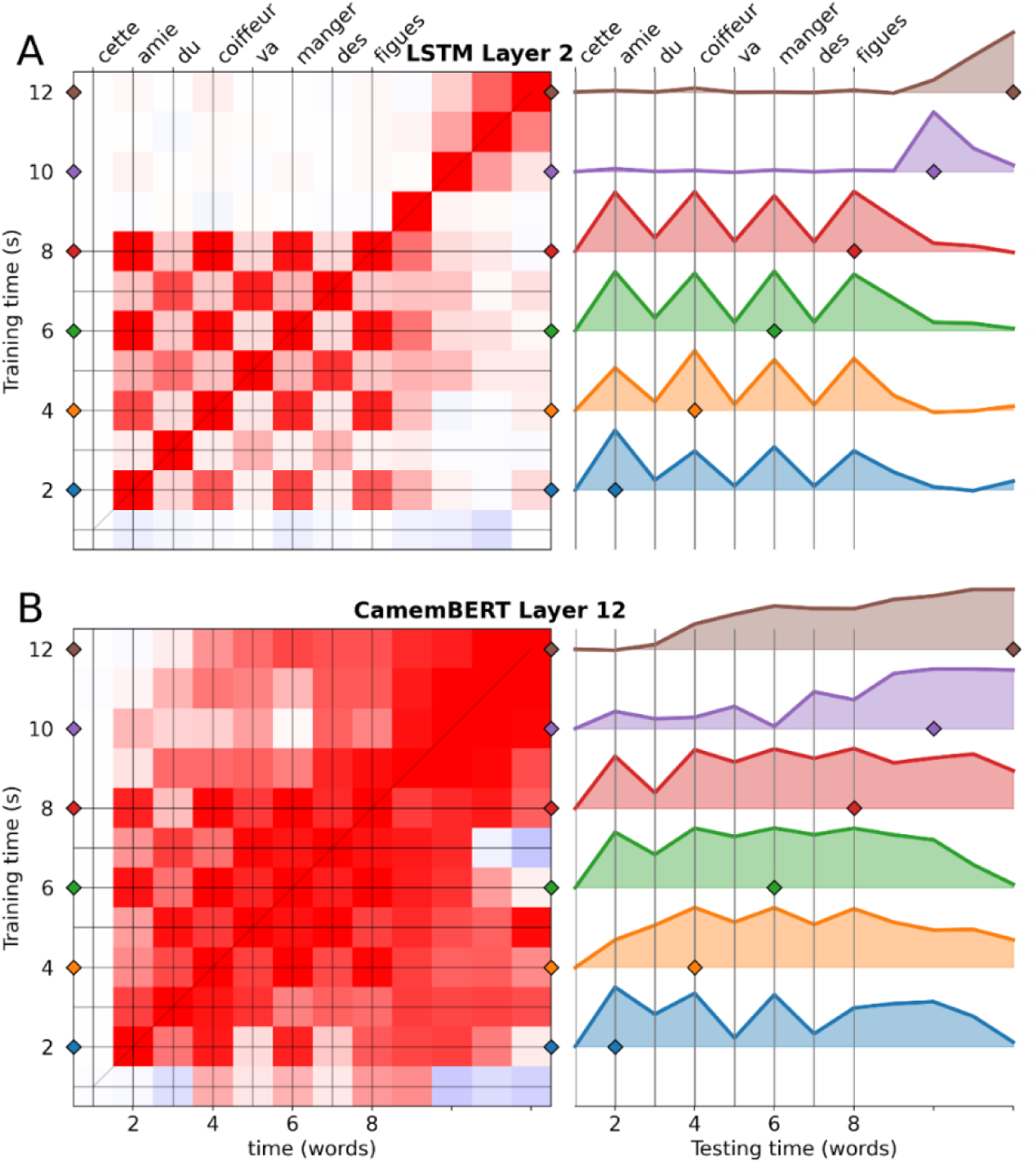
Decoding normal versus Jabberwocky sentences in LSTMs and CamemBERT. Left: Temporal generalization matrices for a decoder trained to distinguish normal sentences from jabberwocky using the activity of an LSTM’s 2nd layer (top) and the CamemBERT’s 12th layer (bottom). Right: Generalization of individual decoders, i.e., horizontal slices from the temporal generalization matrices on the left. These slices from each matrix show in more details the temporal dynamics of sentence processing. Filled lines show significant time points, tested with Wilcoxon-Mann-Whitney test against chance (0.5) and FDR correction.

**Figure 5-2:**
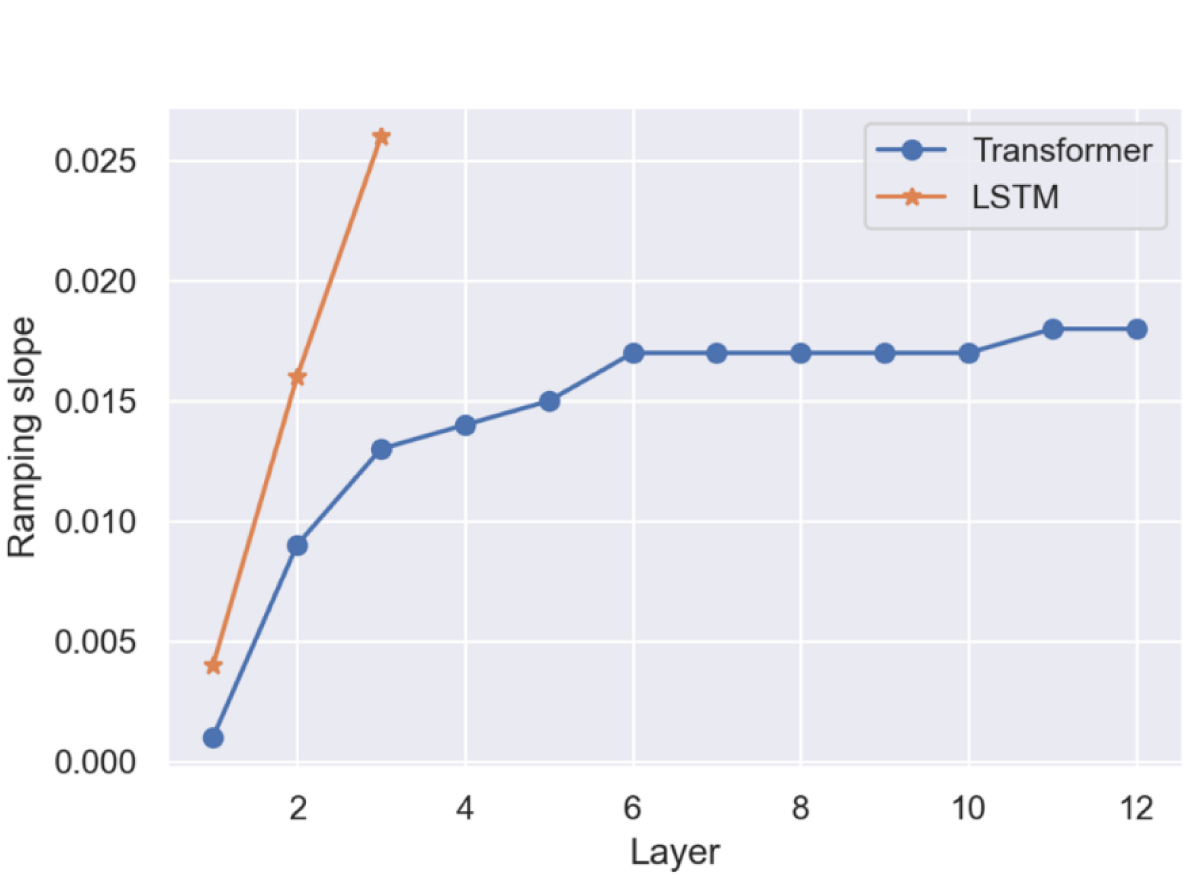
Ramping tendency in LSTMs and Transformers NLMs. Slope of the linear regression on the classification performance between word 1 and 8, for each layer of the LSTMs and Transformers NLMs under study. The ramping slope increases with the layer number. We included all 10 instances of each model in this regression to get substantial statistical power. Because of this, we could not apply this analysis to the CamemBERT model for which a single instance was available.

**Figure 6-1.**
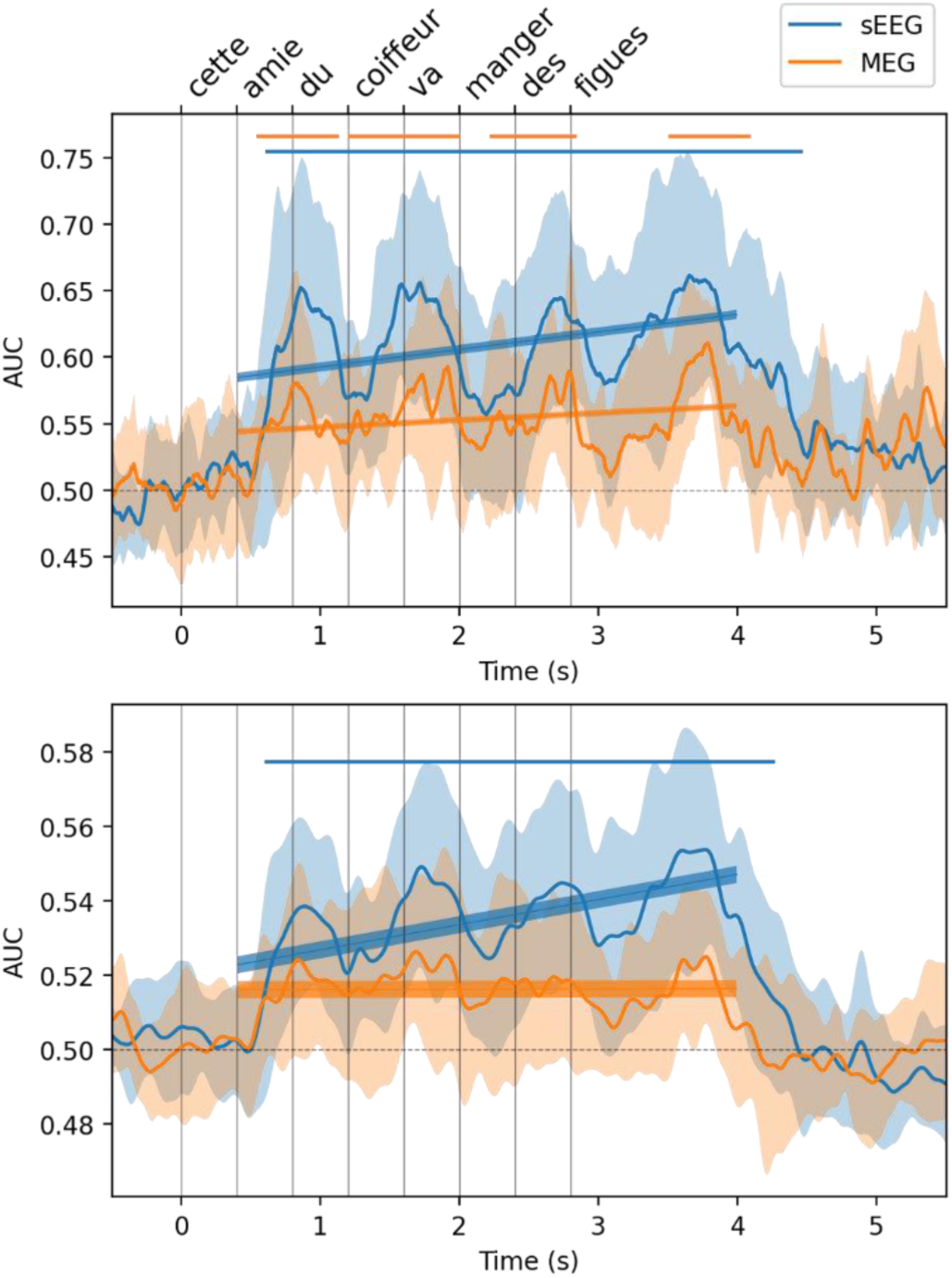
Diagonal decoding performance for normal versus Jabberwocky sentences in human sEEG and MEG. Within-time (diagonal) decoding performance (A) and generalization performance (B), i.e., the average of each line of the temporal generalization matrix. Filled lines show significant time points, tested with cluster permutation test and FDR correction. Regression lines and 95%confidence intervals are also shown. Note that performance stays above chance for a long time after the last word was presented.

**Figure 7-1:**
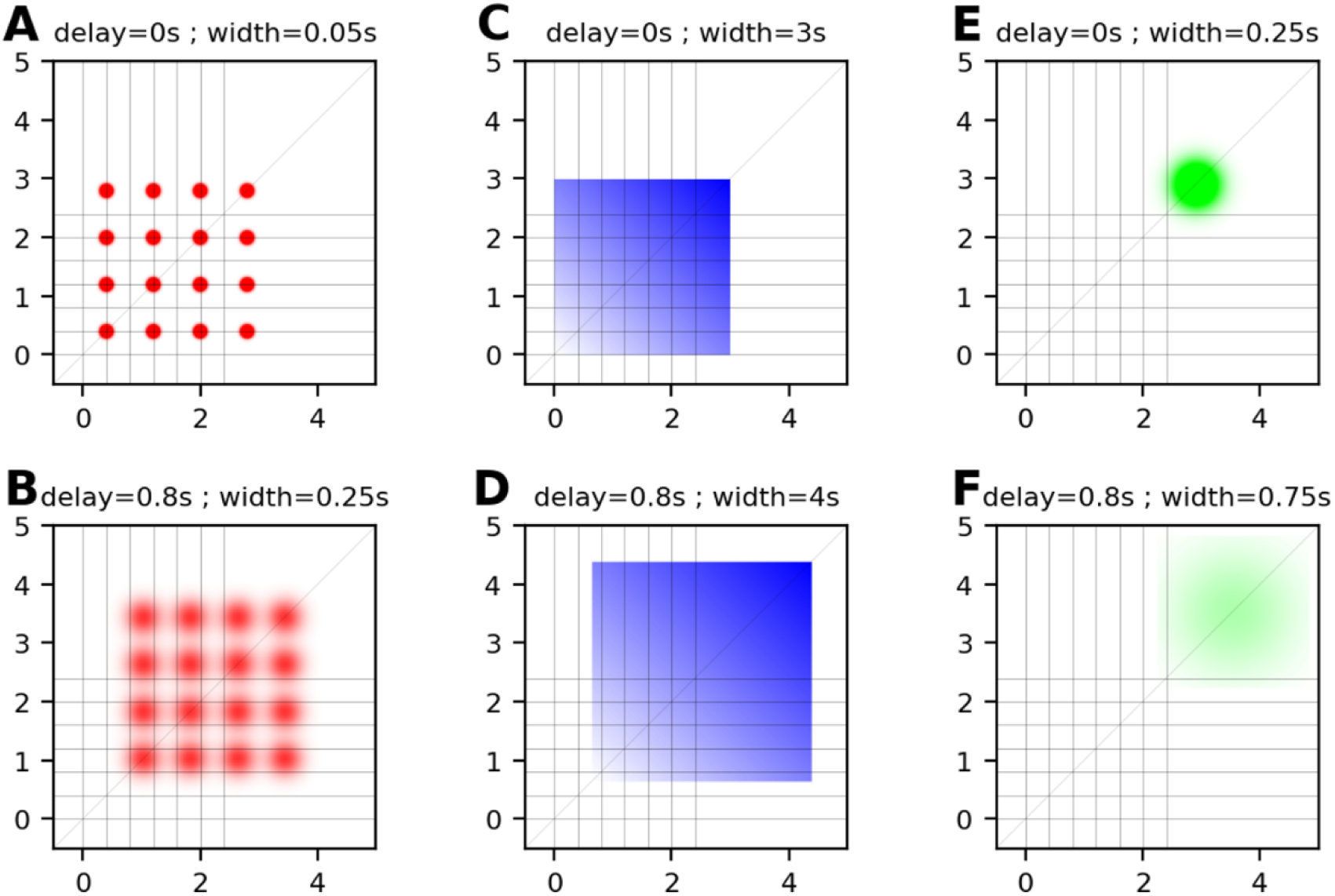
Example templates used in the grid search for the template regression analysis. Matrices shown are the extremes for each parameter (delay and width) selected by the grid search. Each parameter was varied independently of the other. For each template, we tested 10 delays and 10 widths, covering the most likely range of dynamics of interest.

**Figure 7-2:**
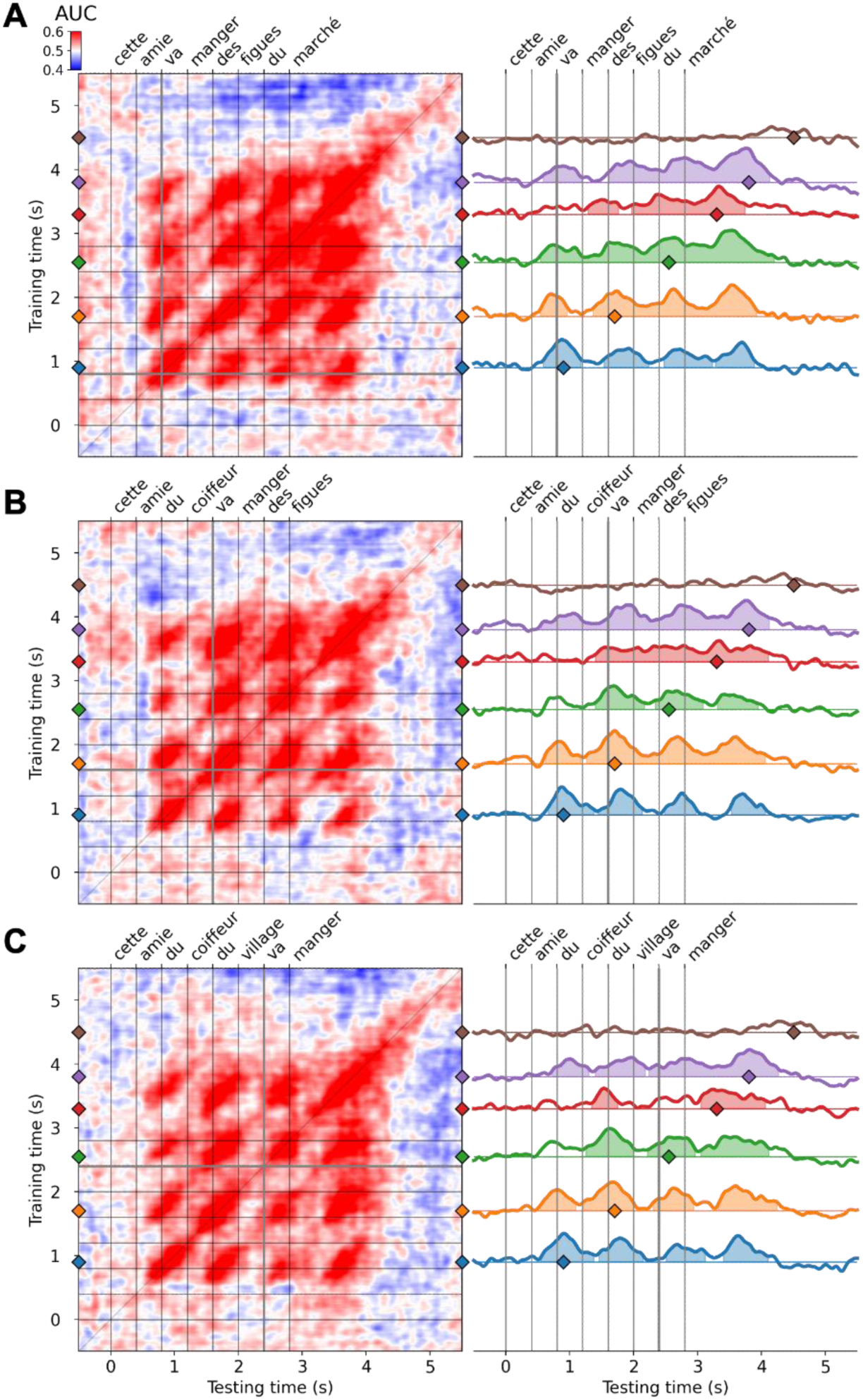
Lack of syntactic modulation of the ramping pattern. The figure shows the temporal generalization matrices of decoders trained on normal versus Jabberwocky using all syntactic structures, and then tested on each structure separately. Matrices correspond to syntactic structure 2-6 (A), 4-4 (B) and 6-2 (C). We tested whether the transition from NP to VP would induce a decrease in decoding performance (peak performance just before the transition, i.e., at the last noun of the NP versus just after the transition, i.e., at the verb, using Wilcoxon-Mann-Whitney tests, but none of these effects were significant. Thick grey lines highlight the transition from NP to VP, where the drop of performance was expected.

